# Constructed languages are processed by the same brain mechanisms as natural languages

**DOI:** 10.1101/2023.07.28.550667

**Authors:** Saima Malik-Moraleda, Maya Taliaferro, Steve Shannon, Niharika Jhingan, Sara Swords, David J. Peterson, Paul Frommer, Marc Okrand, Jessie Sams, Ramsey Cardwell, Cassie Freeman, Evelina Fedorenko

## Abstract

What constitutes a language? Natural languages share features with other domains: from math, to music, to gesture. However, the brain mechanisms that process linguistic input are highly specialized, showing little response to diverse non-linguistic tasks. Here, we examine constructed languages (conlangs) to ask whether they draw on the same neural mechanisms as natural languages, or whether they instead pattern with domains like math and programming languages. Using individual-subject fMRI analyses, we show that understanding conlangs recruits the same brain areas as natural language comprehension. This result holds for Esperanto (n=19 speakers) and four fictional conlangs (Klingon (n=10), Na’vi (n=9), High Valyrian (n=3), and Dothraki (n=3)). These findings suggest that conlangs and natural languages share critical features that allow them to draw on the same representations and computations, implemented in the left-lateralized network of brain areas. The features of conlangs that differentiate them from natural languages—including recent creation by a single individual, often for an esoteric purpose, the small number of speakers, and the fact that these languages are typically learned in adulthood— appear to not be consequential for the reliance on the same cognitive and neural mechanisms. We argue that the critical shared feature of conlangs and natural languages is that they are symbolic systems capable of expressing an open-ended range of meanings about our outer and inner worlds.

**Significance Statement:** What constitutes a *language* has been of interest to diverse disciplines – from philosophy and linguistics to psychology, anthropology, and sociology. An empirical approach is to test whether the system in question recruits the brain system that processes natural languages. In spite of their similarity to natural languages, math and programming languages recruit a distinct brain system. Using fMRI, we test brain responses to stimuli not previously investigated—constructed languages (conlangs)—and find that they are processed by the same brain network as natural languages. Thus, an ability for a symbolic system to express diverse meanings about the world— but not the recency, manner, and purpose of its creation, or a large user base—is a defining characteristic of a language.

## Introduction

What constitutes a ‘language’ has long been debated. Some have discussed this question when comparing human language and non-human animal communication systems (e.g., Hockett, 1959; Premack, 1985; Swadesh, 2017; Call & Tomasello, 2020). Others have argued about whether different human cultural inventions—from math, to music and dance, to cooking, to comics, to programming languages—can or should be considered a kind of language (e.g., McGee, 1997; Hagendoorn, 2010; Patel, 2010; Cohn, 2013; Bers, 2019).

To answer the question of what counts as a ‘language’, one may attempt to provide a set of criteria that must be satisfied for a system to be considered a language. But evaluating any such definition is challenging: one can always argue that the definition should be broader or narrower in scope, include a particular criterion or not, and so on. Alternatively, this question can be answered empirically, by asking whether the cognitive and neural mechanisms that are used to process natural human languages are also used to process information in other domains, like math or music. If so, then presumably, some aspects of the knowledge representations or processing algorithms are sufficiently similar to draw on the same neural circuits.

Over the last few decades, brain imaging studies and investigations of patients with brain damage have revealed that the same brain areas process diverse natural languages, but these areas are strongly selective for language relative to non-linguistic inputs and tasks (Fedorenko et al., 2024a). In particular, the same set of frontal and temporal areas (lateralized to the left hemisphere in the vast majority of individuals; Lipkin et al., 2022) are engaged when native or proficient speakers of Mandarin, Turkish, Pirahã, or any other language tested to date, understand or produce spoken and written linguistic messages (e.g., Illes et al., 1999; Chee et al., 1999, 2000; Roux & Trétmoulet, 2002; Klein et al., 2006; Sulpizio et al., 2020; Malik-Moraleda, Ayyash et al., 2022). The same areas also support the processing of signed languages, which use the visual-manual modality (e.g., Neville et al., 1998; Emmorey et al., 2002; Newman et al., 2015), and whistled languages (Carreiras et al., 2005). These findings hold both when comparing speakers of different languages, and when examining responses to different languages in the brains of multilinguals (e.g., Illes et al., 1999; Vingerhoets et al., 2003; Lemhöfer et al., 2010; De Bruin et al., 2014; Malik-Moraleda, Jouravlev et al., 2023).

In stark contrast to their ubiquitous engagement during the processing of seemingly any of the diverse human languages, language-responsive brain areas are not engaged when individuals solve math problems, understand computer code, listen to music, or perform a wide variety of other tasks (e.g., Monti et al., 2009, 2012; Fedorenko et al., 2011; Amalric & Dehaene, 2019; Ivanova et al., 2020; Liu et al., 2020; Ivanova et al., 2021; Chen et al., 2023). Mirroring these neuroimaging findings, some patients with severe damage to the language system appear to be able to do math, to create and appreciate music, and to engage in diverse forms of reasoning (e.g., Varley & Siegal, 2000; Varley et al., 2005; Apperly et al., 2007; see Fedorenko & Varley, 2016 and Fedorenko et al., 2024b for reviews). Thus, in spite of conceptual parallels that can be drawn between natural languages and other cultural inventions, the machinery that supports our linguistic abilities is not shared with information processing in other domains.

What are the features that all natural languages have that other domains do not? Let us consider a domain that constitutes perhaps the tightest comparison with natural languages yet engages a distinct brain system (Ivanova et al., 2020; Liu et al., 2020): programming languages. Programming languages are compositional generative systems of symbolic form–meaning mappings that allow for the expression of an open-ended range of complex meanings (for discussions of similarities with natural languages, see Bers, 2019; Fedorenko et al., 2019; Casalnuovo et al., 2020). In spite of deep similarities, programming and natural languages differ in important ways. We highlight four salient differences. First, programming languages are evolutionarily much more recent: the earliest ones were only created in the 1940s (Knuth & Pardo, 1980) and many of the currently popular programming languages, such as Python (van Rossum, 1991), have only existed for 2-3 decades, whereas natural languages emerged as early as a million years ago (e.g., Perreault & Mathew, 2012; Dediu & Levinson, 2013; Barham & Everett, 2021) and many modern languages have existed in their current form for a few centuries. Second, programming languages are typically created by a single individual rather than emerging naturally in social groups, as natural languages do (e.g., Jackendoff, 1999; Kirby et al., 2008; Senghas et al., 2014; Ergin et al., 2020). Third, in contrast to natural languages, learning programming languages requires explicit instruction. Finally, programming and natural languages express different kinds of meanings, plausibly due to the different goals for which these systems were created. Natural languages express meanings that relate to the physical and social worlds: they allow us to refer to objects and agents, their properties, describe the actions they undertake and the interactions among them, and the agents’ (including our own) internal emotional and mental states. In this way, natural languages provide a mechanism for sharing our knowledge and beliefs about the world, as well as our feelings and emotions with each other. Programming languages are much more restricted in the kinds of meanings they can express and bear little relationship to our experiences in the world, including interactions with conspecifics. Instead, they express highly abstract, often relational, concepts and are designed to fulfill a particular cultural need (using machines to improve speed and accuracy on diverse tasks). Could one of these differences explain why processing natural vs. programming languages draw on different brain systems? Explicit instruction can be ruled out as a critical factor: natural languages acquired late in life through explicit instruction (e.g., language classes) engage the same brain areas as one’s native language (e.g., Rüschemeyer et al., 2005; Barbeau et al., 2017; Malik-Moraleda, Jouravlev et al., 2023). The other three differences are viable as hypotheses based on the empirical data currently available, although they differ in their intuitive plausibility. In particular, recency of creation alone does not seem *a priori* likely to drive a system to recruit different brain mechanisms as long as the relevant representations and computations are similar to those of standardly studied natural languages.

To help narrow the possibilities for the critical stimulus features that lead to the engagement of the language brain areas, we examined responses to a class of stimuli that, to our knowledge, have not been previously examined in cognitive neuroscience: invented or constructed languages, or conlangs, like Esperanto and Klingon. Similar to programming languages, conlangs i) are relatively recent cultural inventions (albeit older than programming languages) — with the earliest records of conlangs only dating back to the 12th century—and ii) are typically created by single individuals, and often for esoteric purposes, such as creating an identity for a fictional species (**Table 1**; Okrent, 2010). However, similar to natural languages, conlangs are communication systems that can be used to talk about the outer world (the objects/entities and events) and the inner world (our feelings, thoughts, and attitudes). Moreover, conlangs are created by proficient users of natural language and are therefore typically modeled, in some way, on natural languages, even when trying to create languages with unusual properties. In fact, conlang creators are often individuals with heightened linguistic awareness, obtained through acquisition of multiple natural languages and/or formal linguistic training (Schor, 2016). Familiarity with multiple natural languages allows conlang creators to borrow and combine in creative ways features of different languages to create a novel language. In this way, conlangs would be expected to share numerous phonological, morphological, lexical, and syntactic features with natural languages, beyond the similarities in the expressable meanings.

**Table 1:** Information on the conlangs that are included in the present study.

| Conlang | Inception | Number of users<br>(historically) | Creator | Purpose of creation |
| --- | --- | --- | --- | --- |
| Esperanto | 1887 | ~60,000 | Ludwik Lejzer Zamenhof | Facilitate international communication |
| Klingon | 1985 | ~2,000 | Marc Okrand | Fiction |
| Na'vi | 2009 | ~100 | Paul Frommer | Fiction |
| High Valyrian | 2011 | ~50 | David J. Peterson | Fiction |
| Dothraki | 2011 | ~10 | David J. Peterson | Fiction |

We therefore asked whether the processing of conlangs (by proficient users) would engage the brain areas that process natural languages, or whether conlang processing would instead draw on the brain network that supports the processing of computer languages—the so-called Multiple Demand network (Ivanova et al., 2020; Liu et al., 2020). The latter network supports diverse goal-directed behaviors and has been linked to fluid intelligence and learning of new skills (e.g., Duncan, 2010, 2013; Duncan et al., 2020; Assem et al., 2020). The Multiple Demand network is also the network that supports mathematical reasoning and some forms of logical reasoning (e.g., Monti et al., 2009, 2012; Fedorenko et al., 2013; Amalric & Dehaene, 2019; Liu et al., 2020; Kean et al., 2024). To ensure the generalizability of our results, we tested speakers of five conlangs—Esperanto and four fictional conlangs (Klingon, Na’vi, High Valyrian, and Dothraki)—which differ in the span of time they have been spoken, the size of the community that uses the language, the creator, and the purpose for which they were created (**Table 1**). To foreshadow our results, we found that all five conlangs recruit the same brain areas that support the processing of natural languages.

## Results

### Behavioral

Participants reported a range of proficiency levels in their conlang(s), with most participants self-reporting moderate to high proficiency (73.17% of the participants reported proficiency of 3 and above on the 1-6 CEFR scale; M=3.7, range: 1.2-5.8; **Figure 1a**). Higher proficiency levels were reported for Esperanto (*M=*4.0, range: 1.4-5.8), Klingon (*M=*3.3, range: 1.6-5.4), and Na’vi (*M=*4.2, range: 2.6-5.2), compared to High Valyrian (*M=*1.9, range: 1.2-3.2) and Dothraki (*M=*2.8, range: 1.2-4.0). For the open-ended written production task, participants produced 61.68 words on average and 15.83 words per minute on average (Esperanto: *M=*19.26, range: 4.17-44.29; Klingon: *M=*9.26, range: 1.60-23.37; Na’vi: *M=*15.82, range: 2.60-29.78; Dothraki: *M=*9.50, range: 4.73-15.75; High Valyrian was excluded from these analyses because participants in this group did not consider themselves proficient enough to perform the task). And for the lexical decision task, the average accuracy was 0.90 for Esperanto speakers and 0.90 for Klingon speakers (both reliably greater than chance; ts>7, ps<0.001). Furthermore, across the full set of participants, the self-reported proficiency scores were predictive of both the words per minute scores on the production task (r=0.53; t(36)=3.73, p<0.001) and the lexical decision accuracies (r=0.50; t(23)=2.76, p=0.01; **Figure 1b**).

**Figure 1:**
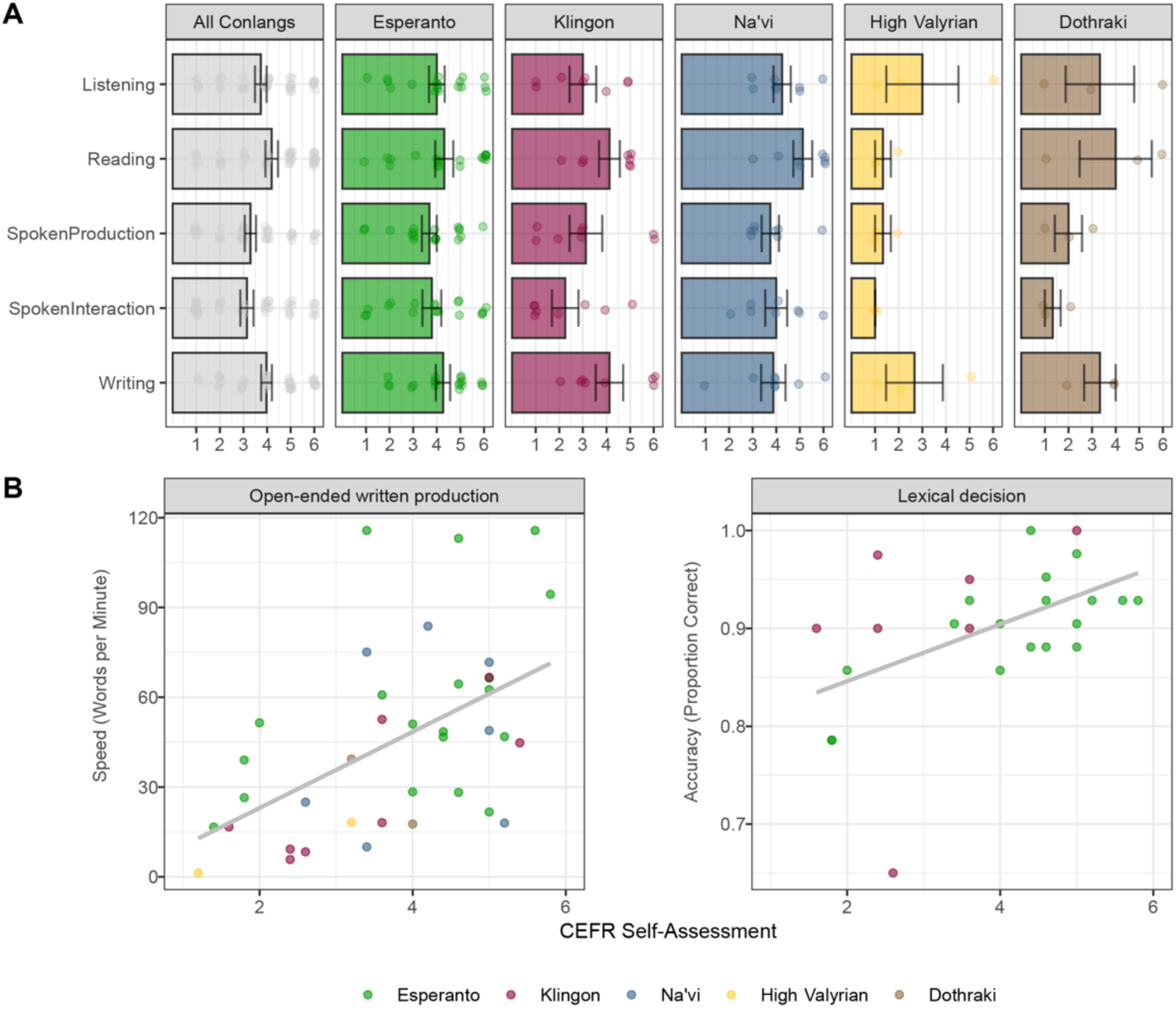
Self-reported proficiencies are in the moderate to high range for most participants and are predictive of open-ended written production ability and lexical decision task accuracies. A) The average self-report CEFR scores for each conlang and across conlangs in the five domains (listening, reading, spoken interaction, spoken production, and writing). Dots represent individual participant CEFR scores. B) Correlation of overall CEFR scores (averaged across the five domains) to the open-ended written production task and the lexical decision task. Dots represent individual participants and are colored according to the conlang.

### fMRI

In line with past results, the language fROIs (defined by the Sentences > Control contrast from the reading-based English localizer) responded more strongly during the *Sentences* condition than the *Control* (nonwords) condition (2.16% vs 0.77% BOLD signal change relative to the fixation baseline, estimated using an independent portion of the data; β=1.40, p<0.001; see **Supp. Table 5** for responses in each conlang group separately). The language fROIs also responded more strongly during the *Sentences* than the *Control* condition in the listening-based localizer administered in the participant’s native language (2.02 vs. 0.59% BOLD signal change relative to fixation; β=1.43, p<0.001; **Supp. Table 5**).

Critically, the language fROIs showed a robust response during the comprehension of constructed languages (**Figures 2** and **3**; see **Supp. Figure 2** for responses broken down by fROI and **Supp. Figure 3** for a comparison of Esperanto vs. the rest of fictional languages). The *Sentences* condition elicited a stronger response than the *Control* condition across conlangs (1.96 vs. 0.75% BOLD signal change relative to fixation; β=1.22, p<0.001). This critical result held for each of the five conlangs individually (**Figure 1**): Esperanto (2.16 vs. 0.89; β=1.27, p<0.001), Klingon (1.36 vs. 0.41; β=0.96, p<0.001), Na’vi (2.67 vs 1.07; β=1.61, p<0.001), High Valyrian (1.37 vs. 0.45; β=0.92, p<0.001) and Dothraki (1.86 vs. 0.73, β=1.13, p<0.001).

**Figure 2.**
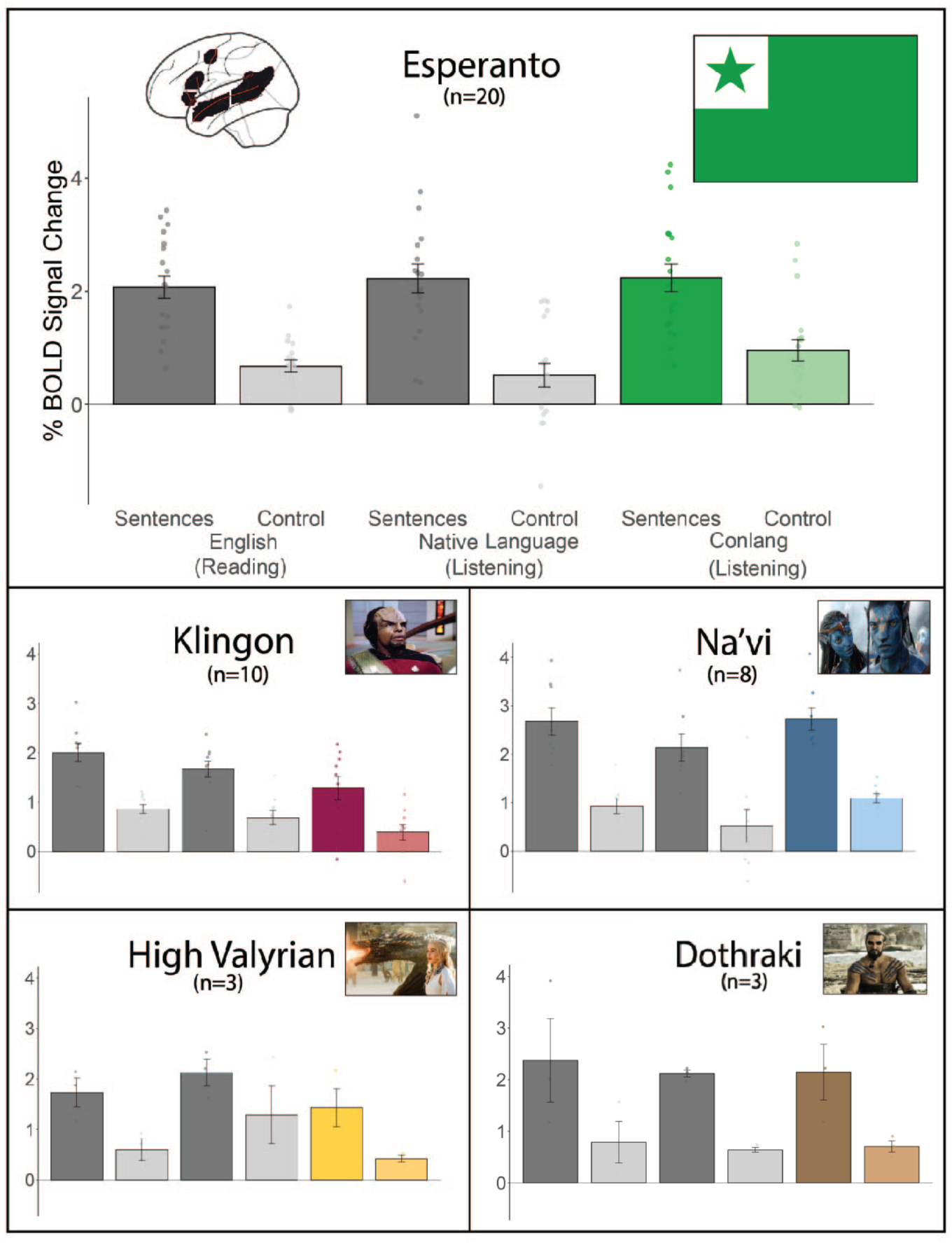
The language areas show a strong response during the processing of both natural and constructed languages. Percent BOLD signal change across the five language fROIs (the results are similar in each fROI separately; **Supp. Figure 2**) for the reading-based and the listening-based versions of the language localizer (dark grey bars = sentences, light grey bars = control condition; the reading-based version used English materials, in which all participants were native or proficient, and the listening-based version used the participants’ native language materials) and the critical conlang comprehension task, where participants listened to sentences in the constructed language (colored bars) and to control, acoustically degraded versions of those sentences (lighter versions of the colored bars). The language network is defined by the Sentences>Control condition of the English reading task in the language-dominant hemisphere (left hemisphere for 36 of the 38 participants), but the results are similar when the language network is defined by the Sentences>Control contrast of the listening language localizer task (**Supp. Figure 1**) or when the left-hemisphere fROIs are used for all participants (**Supp. Figure 5**). Dots represent individual participants; error bars represent standard errors of the mean by participant.

**Figure 3.**
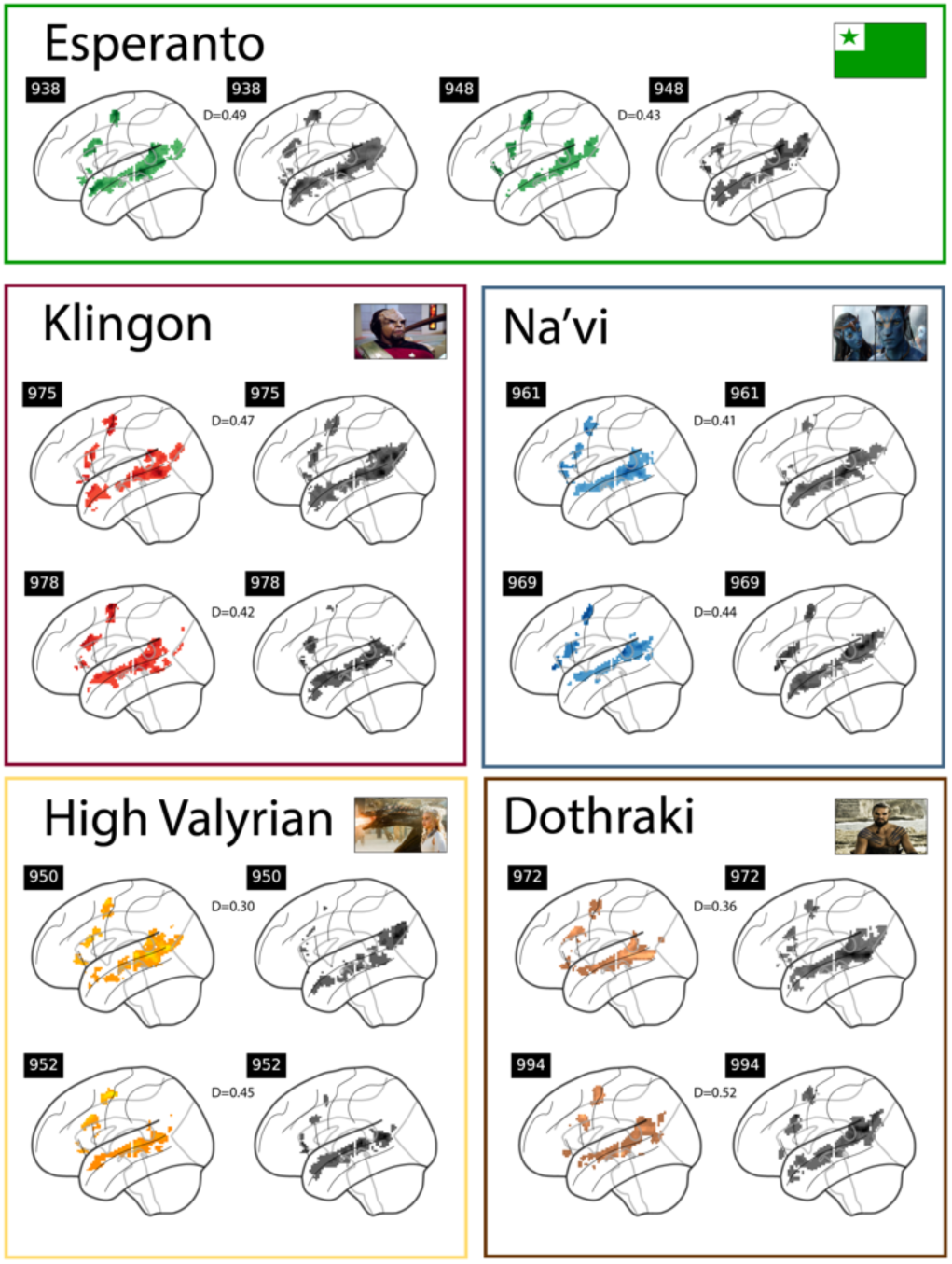
The topography of the language network’s response is similar between constructed languages (different colors, left map for each participant) and natural languages (grey, right map for each participant). For each conlang, we show two sample participants (participant ID—according to a lab-internal numbering system—is shown in small black boxes; these can be cross-referenced with the data tables on OSF). Within each parcel of the language network and for each language, the 10% of most language-responsive voxels are shown. The D values correspond to the Dice coefficient value (computed across the brain, as described in the text) for the comparison between the activations in response to the participant’s native language vs. their conlang. All the individual maps are available at https://osf.io/pe74v/.

To further characterize the similarity of brain responses to natural vs. constructed languages, we created binarized versions of the activation maps for the Sentences>Control contrast for the participant’s native language and the relevant conlang where—restricting the analysis to the LH language parcels—voxels that belonged to the set of 10% of most responsive voxels were assigned a value of 1 and voxels that did not were assigned a value of 0 (**Figure 3; Supp. Figure 4** shows the results when selecting the top 5% voxels and top 20% of voxels instead). We then computed the Dice coefficient (Rombouts et al., 1997), which ranges from 0 (no overlapping voxels) to 1 (all voxels overlap) between a) the native language map and the conlang map within each participant (n=38 pairs), and b) the native language maps between each pair of participants (n=703 pairs). We found that the sets of most responsive voxels are more similar (show greater overlap) within an individual between a natural and a constructed language (0.45) than between individuals for a natural language (0.31; β=0.23, p<0.001; see **Figure 3** for sample maps). This result further supports the claim that conlangs are represented and processed in a similar brain network to a native language within individual speakers.

As discussed in the Introduction, language areas have been shown to be selective for language processing relative to diverse cognitive tasks (e.g., Fedorenko et al., 2011; reviewed in Fedorenko et al., 2024a). Here, we replicate this finding in this new set of participants with respect to two non-linguistic tasks and directly compare the magnitude of the response for conlang comprehension to the magnitude for these non-linguistic conditions. As can be seen in **Figure 4**, the response in the language fROIs to the four non-linguistic conditions (a hard and an easy condition of a spatial working memory task and a mental and a physical condition of a naturalistic Theory of Mind paradigm) falls at or below the fixation baseline (in line with e.g., Mineroff, Blank et al., 2018; Malik-Moraleda, Ayyash et al., 2022; Shain, Paunov, Chen et al., 2022). The response to the sentence condition in the conlang experiment is comparable to the magnitude of response to the sentence condition in the two language localizer paradigms and is reliably higher than each of the four non-linguistic conditions (ps<0.001; see **Supp. Table 5** for the results broken down by conlang; see **Supp. Figure 6** for evidence that the non-linguistic paradigms elicit the expected response in the Multiple Demand and Theory of Mind areas; see **Supp. Figure 7** for the whole-brain overlap analysis between conlang activations and the language localizer activations vs. non-linguistic task activations).

**Figure 4.**
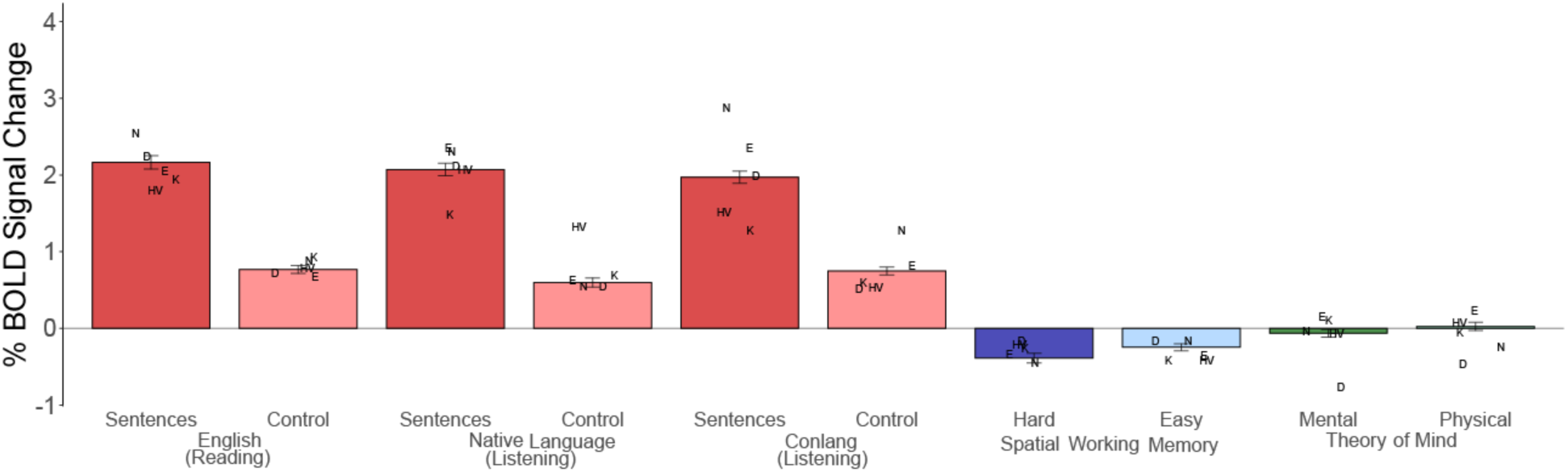
The language areas of conlang speakers are selective for language: the response to conlangs is similar in magnitude to the response to natural languages and reliably higher than the response to non-linguistic tasks. The letters correspond to the average response for each constructed language (E: Esperanto; K: Klingon; N: Na’vi; HV: High Valyrian; D: Dothraki).

## Discussion

In an fMRI investigation of neural responses to constructed languages (conlangs), we found that conlangs recruit the same brain areas—frontal and temporal areas in the left hemisphere—as those that support the processing of natural languages. This result held for both Esperanto and fictional languages. Esperanto is a conlang that was created over 130 years ago to facilitate international communication, was explicitly modeled on natural languages, and has a substantial number of speakers. In contrast, the four fictional conlangs (Klingon, Na’vi, Dothraki, and High Valyrian) were all created within the last 12-40 years (**Table 1**), were constructed so as to differ from most natural languages by using linguistic properties that are uncommon across the world’s languages (two of them were created for fictional humanoid alien species), and have relatively small communities of speakers. Although the five conlangs differ in the strength of the response they elicit in the language areas (with Esperanto and Na’vi eliciting higher responses than Klingon, High Valyrian, and Dothraki), these differences are likely due to differences in the proficiency levels among the speakers (e.g., see Malik-Moraleda, Jouravlev et al., 2023 for evidence that the level of response in the language areas scales with proficiency). Below we discuss some implications of these findings for understanding the human language system.

We started this paper with a discussion of how to determine whether some information system can/should be considered a ‘language’. We advocated an empirical approach: testing whether some system of interest engages the same neural machinery as natural languages. If so, then we can infer that the system in question shares some aspects of representation and/or computation with natural languages. Past work has shown that the language brain areas are similar across speakers of typologically diverse languages (e.g., Malik-Moraleda, Ayyash et al., 2022) but are not engaged by diverse non-linguistic stimuli that share features with natural languages (see Fedorenko et al., 2024a for review). Focusing on programming languages—which share deep similarities with natural languages (Fedorenko et al., 2019)—we discussed three features that could explain why the processing of programming languages draws on a distinct brain system: i) their evolutionary recency, ii) their mode of creation (intentionally, by a single individual), and iii) the kinds of meanings they express. Conlangs share the first two features with programming languages and the third feature—with natural languages. The fact that conlang comprehension recruits the same brain areas as natural language processing rules out ancient evolutionary history and natural emergence in social groups as critical features of systems that engage the language-processing mechanisms. In this way, our work extends prior research on the ubiquity of engagement of the language areas by diverse languages, including spoken languages across language families (e.g., Malik-Moraleda, Ayyash et al., 2022), signed languages (e.g., Neville et al., 1998; Emmorey et al., 2002; Newman et al., 2015), and whistled languages (Carreiras et al., 2005). In addition to the features mentioned above, our findings suggest that to elicit a robust response in the language areas, a language does not need to be used by an extensive community of speakers, and it does not need to have been shaped by learning pressures, at least not by the kinds of pressures that act on first language acquisition. It is important to acknowledge, however, that we have only focused on a small number of conlangs, all of which ended up being learnable and used by a community of speakers. At the same time, conlangs are being created daily all over the world, from languages created by twins or close siblings (e.g., Bakker, 1987; Mogford-Bevan, 1999; Thorpe, 2006) to languages created by language aficionados and never shared with anyone, to languages created so as to explicitly remove some key features of natural language, such as ambiguity (e.g., Lojban; 2023), including languages that prove difficult to learn (e.g., Ithkuil; Quijada, 2011). Whether all of these conlangs would engage language-processing mechanisms in their learners and users is not yet known.

On the other hand, the fact that conlangs, but not programming languages (or mathematical and logic expressions; Monti et al., 2009, 2012; Fedorenko et al., 2011; Kean et al., 2024), engage the language areas suggests that the nature of the meanings conveyed by these different symbolic systems matters for which brain system is engaged. Note that it is not the abstract underlying meanings per se that matter: non-verbal stimuli matched for meaning elicit a relatively weak response in the language areas (e.g., Ivanova et al., 2021). The critical feature appears to be the symbolic forms that lead to the activation of the associated meaning structures.

Might something about the form-related features themselves be important? As noted in the Introduction, conlangs and natural languages are similar not only in the kinds of meanings they can express, but also in their formal features, including how words are formed and how they are combined into phrases and sentences. The importance of particular form-based features for engaging the language system is less clear. One past fMRI study tested whether the learning of real vs. made-up syntactic rules engages language-processing mechanisms (Musso et al., 2003). The researchers used natural languages and taught participants who were unfamiliar with those languages a small vocabulary and either real rules for combining the words or made-up rules (which violate a particular syntactic theory’s constraints). Based on observing activation within the left inferior frontal gyrus for only the real rules, they argued that only the processing of real rules, allowed by the grammars of natural languages, engages the language areas. However, because the language areas were not functionally defined in that study, it is difficult to know whether the observed effects localize to the language cortex or the nearby Multiple Demand cortex, which responds to diverse demanding tasks (Fedorenko et al., 2012; see Fedorenko & Blank, 2020 for discussion).

Other studies have investigated the importance of syntactic features for engaging the language areas by probing neural responses to linguistically degraded conditions, which lack most of linguistic meaning, but preserve the grammatical frame: so-called “Jabberwocky” sentences (Carroll, 1871). Such studies have found that responses of the language areas to such stimuli are substantially lower than to natural language sentences (Fedorenko et al., 2010; Shain, Kean et al., 2024). Yet others have taken inspiration from classic statistical learning studies by Saffran and colleagues (Saffran et al., 1996; for a review, see Saffran & Kirkham, 2018) and used auditory (or, sometimes, visual) sequences with no meanings associated with any elements or element combinations. These paradigms focus on the learning of structural regularities (e.g., a particular syllable always follows some other syllable or syllable type, or a particular pattern repeats, like A B B A). For these paradigms, some have reported responses in putative language areas (e.g., Bahlmann et al., 2008; Petersson et al., 2012; Uddén & Bahlmann, 2012; Wang et al., 2015; Friederici et al., 2006; see e.g., Uddén & Männel, 2018 for a review). However, when the language areas are defined functionally and thus the observed activations can be unambiguously interpreted as arising within the language cortex (Fedorenko et al., 2012; see Fedorenko & Blank, 2020 for discussion), learning statistical regularities in sequences of meaningless elements elicits weak or no response in the language areas (Schneider et al., 2024). Thus, form-based regularities on their own are not sufficient to engage the these areas. The relatively low responses in the language areas to Jabberwocky sentences further shows that even the formal features of a familiar language (e.g., grammatical morphological endings and function words) only engage the language system weakly, presumably because such stimuli do not allow for the full range of linguistic computations, like those related to lexical access and syntactic structure building (see Malik-Moraleda, Jouravlev et al., 2023, for a related discussion in the context of processing low-proficiency languages). But to understand whether particular formal properties that characterize natural and constructed languages are critical for engaging the language mechanisms would require additional investigation.

For example, is compositionality required for a communication system to engage the language brain areas? To the best of our knowledge, no cognitive neuroscience research has been carried out on the processing of holistic languages (e.g., Wray, 2000)—languages made up of random pairings of signals and meanings. A fruitful paradigm for experimentally evaluating the importance of compositionality and other features of communication systems for engaging language-processing mechanisms might be the iterated learning paradigm introduced to the domain of communication by Kirby and colleagues (e.g., Smith et al., 2003; Kirby et al., 2008, 2014, 2015; Motamedi et al., 2019; see Kirby, 2017 for an overview). In contrast to artificial grammar learning paradigms (e.g., Uddén & Männel, 2018), where the sequences are meaningless and therefore do not strongly engage the language mechanisms, iterated language learning paradigms have a meaning component: participants learn labels for some structured set of meanings (e.g., moving objects that vary in shape and type of motion) and transmit these labels to the next generation of learners. Under these conditions, within a couple of ‘generations’ of learners, the initially introduced idiosyncratic labels become systematic and compositional, mirroring the structure of the meaning space (e.g., the first syllable may end up referring to the shape and the second—to the type of motion). Probing brain responses of participants in different generations (before and after a certain feature of the communication system has emerged) could illuminate the importance of different features for engaging language-processing mechanisms.

Another approach is to study natural communication systems that are not considered full-blown languages, including pidgins—communication systems that are created when two or more groups of speakers of different languages are brought together and need to interact (e.g., Bickerton, 1976; Muysken & Smith, 1995; Blasi et al., 2017), homesign systems used by deaf children for communicating with hearing caretakers in the absence of access to sign language (e.g., Goldin-Meadow, 1993; Brentari & Coppola, 2013), and emergent languages in the early stages of creation (e.g., Senghas et al., 2014; Ergin et al., 2020). Such systems can express quite a wide array of meanings but lack some of the structure and systematicity that characterize fully developed languages. If structure and systematicity are critical for engaging the language processing mechanisms, then such systems may be expected to not draw on the language areas or to elicit a weaker response. On the other hand, if flexible expression of complex meanings is the driving force, they should draw on the language system and elicit as strong a response as full-blown natural languages. One difficulty here is to develop ways to robustly quantify the degree of (different aspects of) systematicity in these different communication systems.

Yet another, more challenging, approach to investigating how particular features of language modulate the level of response in the language areas is to leverage variability across natural languages along various dimensions (e.g., some languages have stricter word orders than others, Levshina et al., 2023). The challenge has to do with matching or controlling for all other features of the experimental materials and matching participant groups, or testing large enough groups so that the estimate approaches the population mean (see Malik-Moraleda, Ayyash et al., 2022 for discussion). A related approach is within-language comparisons. For example, Arnon et al. (2010) showed sensitivity in behavioral measures to the frequency of four-word sequences (e.g., “don’t have to worry” vs. “don’t have to wait”, where all the individual words, bi-grams, and tri-grams are matched in frequency, but the former sequence of four words is more frequent), which suggests that we store such chunks as units. Would the processing of these stored chunks require less neural activity than sequences that require composition? We don’t yet know (cf. evidence from processing of morphologically complex vs. simple forms; Bozic et al., 2010).

To conclude, the human brain contains a network of areas that are highly selective for the processing of language: they are recruited when we understand and produce language (e.g., Menenti et al., 2010; Silbert et al., 2014; Hu, Small et al., 2022; Zada et al., 2024) but show little or no response when we engage in diverse non-linguistic behaviors, from processing math and computer code, to music perception, to interpreting facial expressions and gestures, to reasoning about others’ minds (e.g., see Fedorenko & Varley, 2016 and Fedorenko et al., 2024a for reviews). Here, we evaluated brain responses to five constructed languages and showed that they engage the same brain areas as those that support the processing of natural languages and to a similar extent, despite several notable differences between conlangs and natural languages. This work suggests that conlangs share some critical features of natural languages, including the ability to express diverse meanings about external and internal worlds and some form-based features. By testing neural responses to a wider array of naturally emerging and artificially constructed communication systems, future work should continue to distill the necessary and sufficient conditions for being a language according to this empirical criterion.

## Methods

### General approach

Each participant—all proficient speakers of one or more conlangs (see <u>Participants</u> for details)—performed several tasks in an fMRI scanner. The general approach we adopt relies on functionally identifying areas of interest in each individual participant (e.g., Saxe et al., 2006; Fedorenko et al., 2010). This approach has been shown to confer greater sensitivity, functional resolution, and interpretability than the traditional group-averaging fMRI approach (e.g., Saxe et al., 2006; Nieto-Castañón & Fedorenko, 2012; Fedorenko, 2021). As detailed below, each participant performed a) two ‘localizer’ tasks for the language network: a reading-based version (reading English sentences vs. nonword sequences; Fedorenko et al., 2010) and a listening-based version (listening to short passages in one’s native language vs. to acoustically degraded versions of those passages; Malik-Moraleda, Ayyash et al., 2022); and b) the critical conlang task that was similar to the listening-based localizer but used the materials in the relevant conlang. In addition, participants performed c) two non-linguistic tasks, which were used as additional controls to which neural responses to conlangs could be compared (and could also be used for identifying additional candidate networks that could support conlang comprehension).

In addition to the fMRI component, participants self-assessed their conlang proficiency using the Common European Framework of Reference (CEFR; Council of Europe, 2001) grid and wrote short passages (timed) about three topics in the conlang(s) they were proficient in. Esperanto and Klingon speakers additionally completed a lexical decision task.

### Participants

Speakers of constructed languages (all native or highly proficient speakers of English) were recruited online and through word of mouth. Participants were selected based on a) their self-rated proficiency in the constructed language of interest on an informal scale from 1 (“know a couple words”) to 5 (“can understand almost everything and express myself freely”) (see **Supp. Table 1** for details) and b) geographical proximity to Boston, where the testing took place. Overall, 38 individuals (25 males; mean age = 32.31, SE = 1.83) participated in the study, five of whom spoke and were tested on two constructed languages. The pool of participants included speakers of Esperanto (n = 19, 12 males, mean age = 30.47, SE = 2.07), Klingon (n = 10, 6 males, mean age = 40.30, SE=3.64), Na’vi (n = 8, 5 males, mean age = 28.62, SE = 1.89), High Valyrian (n = 3, 1 male, mean age = 40.67, SE =5.49), and Dothraki (n = 3, 2 males, mean age = 37.00, SE = 3.06). A substantial number of participants (17 of the 38 participants) were polyglots, as defined by having some proficiency in 5 or more languages (e.g., Jouravlev et al., 2019). Most participants were right-handed (n = 33), as determined by the Edinburgh Handedness Inventory (n = 35) or by self-report (n=3), and had normal or corrected-to-normal vision. All participants gave informed written consent in accordance with the requirements of MIT’s Committee on the Use of Human as Experimental Subjects, were paid for their participation and, if coming from outside of the Boston area, reimbursed for travel expenses.

## Experimental tasks

### Language localizers

We included two localizer tasks: a visual (reading-based) version (e.g., Fedorenko et al., 2010), which is most commonly used in the Fedorenko lab (e.g., see Lipkin et al., 2022 for data from >600 participants on this version), and an auditory (listening-based) version. The latter was included because 2 of the participants were not native speakers of English, and although we have previously established that an English-based localizer works well for proficient speakers (Malik-Moraleda, Ayyash et al., 2022), we wanted to additionally include a localizer in their native language, and we only have *auditory* versions of localizers for those languages. In the main analyses, we used the reading-based version to define the language areas, and then examined the responses to the conditions of both localizers, to the conditions of the critical task, and the non-linguistic tasks (see **Supp. Figure 1** for evidence that the results are similar when the listening-based version of the localizer is used, as has been previously established; e.g., Fedorenko et al., 2010; Malik-Moraleda, Ayyash et al., 2022; Chen et al., 2023).

#### Reading-based language localizer task

Participants passively read English sentences and lists of pronounceable nonwords (the control condition) in a blocked design. The Sentences > Control contrast targets brain regions that support high-level linguistic processing. Each trial started with 100-ms pre-trial fixation, followed by a 12-word-long sentence or a list of 12 nonwords presented on the screen one word/nonword at a time at the rate of 450 ms per word/nonword. Then, a line drawing of a finger pressing a button appeared for 400 ms, and participants were instructed to press a button whenever they saw this icon. Finally, a blank screen was shown for 100 ms, for a total trial duration of 6 s. The simple button-pressing task was included to help participants stay awake and focused. Each block consisted of three trials and lasted 18 s. Each run consisted of 16 experimental blocks (eight per condition) and five fixation blocks (14 s each), for a total duration of 358 s (5 min 58 s). Each participant performed two runs. Condition order was counterbalanced across runs. This localizer is available from the Fedorenko laboratory website: https://evlab.mit.edu/funcloc/.

#### Listening-based language localizer task

Participants listened to passages from Alice in Wonderland (in their native language; English for n=36, German for n=1, and Dutch for n=1) and to acoustically degraded versions of the passages (the control condition). The acoustically degraded versions were created as described in Scott et al. (2017; see also Malik-Moraleda, Ayyash et al., 2022; **Supp. Text 1**) and sound like poor radio reception such that the linguistic content is not discernible. Similar to the reading-based version, the Sentences > Control contrast targets brain regions that support high-level linguistic processing. Each block (consisting of an intact or a degraded passage) lasted 18 s. Each run consisted of 12 experimental blocks (6 per condition) and 3 fixation blocks (12 s each), for a total duration of 252 s (4 min 12 sec). Each participant performed two runs. Condition order was counterbalanced across runs. The sound was delivered through Sensimetrics earphones (model S14). This localizer is available from the Fedorenko laboratory website: https://evlab.mit.edu/aliceloc.

### Critical conlang task

#### Materials

For each conlang, a set of 100 phrases and sentences was recorded (all the materials are available at: https://osf.io/pe74v/, see **Supp. Table 2** for the English translations). The materials differed across languages, but all sentences were relatively simple, short (**Supp. Table 3**), semantically and syntactically diverse, and—for the fiction conlangs—did not include any show-/movie-specific references. For all conlangs except for High Valyrian, the materials were recorded by a highly proficient male speaker. For Esperanto and Klingon, initial sentence recordings were provided by researchers at Duolingo, but the sentences were then re-recorded by proficient male speakers; for Na’vi, P.F. recorded the sentences; for Dothraki, D.P. recorded the sentences. For High Valyrian, due to the low number of proficient speakers, we used the sentence recordings provided by Duolingo (recorded by several male speakers). Acoustically degraded versions of the materials (for the control condition) were created in the same way as the degraded versions in the auditory version of the language localizer (**Supp. Text 1**). The volume levels were normalized across all clips. The recordings are available at: https://osf.io/pe74v/.

#### Procedure

For each conlang, the sentences were concatenated in order to create 16 blocks of similar length; a similar procedure was applied to the acoustically degraded versions of the sentences. Each block consisted of 5-8 sentences/degraded sentences (see **Supp. Table 3** for details by language), with a 0.5 second pause between sentences, and was between 20 and 24 seconds in duration depending on the conlang (**Supp. Table 3**). For blocks that were a little longer than the target length, a gradual volume fade-out was applied during the last 2 seconds of the block. The 32 blocks (16 Sentences, 16 Control) were further divided into two sets corresponding to two scanning runs, with each run containing 16 blocks (8 Sentences, 8 Control) as well as five 12 seconds-long fixation blocks and lasting between ∼6 and 7.5 minutes depending on the conlang (**Supp. Table 3**). Participants were asked to listen attentively. (For Dothraki, a subset of the materials was discovered during the data analysis stage to not be proper Dothraki (**Supp. Table 2**). These materials affected 5 of the 16 blocks. We modeled those 5 blocks as a separate condition of no interest, so for Dothraki, the critical condition only consists of 11 blocks.)

### Non-linguistic tasks

We included two non-linguistic tasks—a spatial working memory task and a naturalistic Theory of Mind task—which were used as additional controls to which neural responses to conlangs could be compared. The spatial working memory task (Fedorenko et al., 2013) also served as a localizer for the Multiple Demand network (Duncan, 2010, 2013)—the network that has been found to support comprehension of programming languages (e.g., Ivanova et al., 2020) and that served as the main alternative hypothesis for which brain system conlang comprehension would recruit.

#### Spatial Working Memory task

Participants had to keep track of four (easy condition) or eight (hard condition) locations in a 3 × 4 grid. In both conditions, participants performed a two-alternative forced-choice task at the end of each trial to indicate the set of locations that they just saw. Each trial lasted 8 s (see Fedorenko et al., 2011 for the trial timing details). Each block consisted of four trials and lasted 32 s. Each run consisted of 12 experimental blocks (six per condition) and four fixation blocks (16 s in duration each), for a total duration of 448 s (7 min, 28 s). Each participant performed two runs. Condition order was counterbalanced across runs. The hard > easy contrast targets brain regions engaged in diverse demanding cognitive tasks (e.g., Fedorenko et al., 2013; Shashidhara et al., 2019; Assem et al., 2020). This localizer is available from the Fedorenko laboratory website (https://evlab.mit.edu/funcloc/).

#### Naturalistic Theory of Mind task

Participants watched a short silent animated film ("Partly Cloudy", Pixar Animation Studios; 5 min 36 s in duration). The film has been coded for four types of events: i) mental (events that elicit thoughts about a character’s mental state; e.g., the cloud character believing that his friend has left him; 4 events, 44 s total); ii) social (events that involve social engagement but no specific mental/emotional content; e.g., the cloud character and a stork playing together; 5 events, 28 s total); iii) physical pain (e.g., the stork being bitten by a crocodile; 7 events, 26 s total); and iv) control (events where no characters are involved: e.g., shots of the landscape; 3 events, 24 s total). The mental > physical contrast targets brain regions engaged in thinking about another person’s thoughts and beliefs (Jacoby et al., 2016; Richardson & Saxe, 2020; Shain, Paunov, Chen et al., 2022).

### Behavioral tasks

#### CEFR self-assessment grid

Participants were asked to rate their proficiency in the relevant conlang(s) using the CEFR self-assessment grid developed by the Council of Europe (2001). In particular, they were asked to rate their proficiency in each of five different modalities: listening, reading, spoken interaction, spoken production, and writing. For each modality, participants were presented with six statements that describe basic to advanced skills and had to select the sentence that best describes their ability in the relevant conlang. For example, for listening, basic skills are described by the following statement: “I can understand familiar words and very basic phrases concerning myself, my family and immediate concrete surroundings when people speak slowly and clearly.”; and advanced skills are described by: “I have no difficulty in understanding any kind of spoken language, whether live or broadcast, even when delivered at fast native speed, provided I have some time to get familiar with the accent.” These statements correspond to the set of six common reference levels (A1, A2, B1, B2, C1, C2) developed by the Council of Europe to describe proficiency in a language (see **Supp. Table 4** for details on each reference level). For each participant, the scores were averaged across the five modalities to derive a single score (range: 1 (corresponding to A1) to 6 (corresponding to C2)).

#### Open-ended written production task

Participants were asked to write a few sentences about three topics in the conlang of interest. For each participant, the topics were randomly selected from the following set: family, hobbies, animals, home, friends, food, weather, sports, and clothes. On each trial, the topic was presented at the top of the screen along with the instructions below it (“please write about the topic above”). Participants had to type their responses into a text box window and then click the “submit” button at the bottom of the screen once they finished typing. The responses were timed from the first click on the text box until the “submit” button was pressed. For each participant, we calculated the average number of words per minute for each topic and averaged these values across the three topics to derive a single score for each participant.

#### Lexical decision task

Speakers of Esperanto and Klingon were presented with 40 strings of letters, one at a time, and were asked to decide as quickly as possible whether each string was a real word in the relevant conlang. Twenty of these strings were real words and 20 were nonwords. To create the materials, we started with a set of 40 English words from the set of words used in the Peabody Picture Vocabulary Test (PPVT; Dunn & Dunn, 1965) that exist in both Esperanto and Klingon. Twenty of these words were translated into Esperanto / Klingon and used as real words. The other twenty words were translated and then altered (by removing, adding, or transposing some of the letters) and cross-referenced with conlang dictionaries to ensure that the resulting strings did not constitute real words. The order of strings was randomized for each participant. On each trial, the letter string was presented in lower-case letters in the center of the screen, and participants were asked to press one of two buttons on the keyboard to indicate their response. For each participant, we calculated the accuracy score (number of correctly answered trials out of 40). Reaction times were recorded but not used in the analyses.

### fMRI data acquisition

Structural and functional data were collected on a whole-body 3 Tesla Siemens Prisma scanner with a 32-channel head coil at the Athinoula A. Martinos Imaging Center at the McGovern Institute for Brain Research at MIT. T1-weighted, Magnetization Prepared Rapid Gradient Echo (MP-RAGE) structural images were collected in 208 sagittal slices with 1 mm isotropic voxels (TR = 1,800 ms, TE = 2.37 ms, TI = 900 ms, flip = 8 degrees). Functional, blood oxygenation level-dependent (BOLD) data were acquired using an SMS EPI sequence with a 90° flip angle and using a slice acceleration factor of 3, with the following acquisition parameters: seventy-two 2 mm thick near-axial slices acquired in the interleaved order (with 10% distance factor), 2 mm × 2 mm in-plane resolution, FoV in the phase encoding (A >> P) direction 208 mm and matrix size 104 × 104, TR = 2,000 ms, TE = 30 ms, and partial Fourier of 7/8. The first 10 s of each run were excluded to allow for steady state magnetization.

### fMRI data pre-processing and first-level modeling

fMRI data were analyzed using SPM12 (release 7487), CONN EvLab module (release 19b) and other custom MATLAB scripts. Each participant’s functional and structural data were converted from DICOM to NIFTI format. All functional scans were co-registered and resampled using B-spline interpolation to the first scan of the first session. Potential outlier scans were identified from the resulting subject-motion estimates as well as from BOLD signal indicators using default thresholds in CONN pre-processing pipeline (5 s.d. above the mean in global BOLD signal change or framewise displacement values above 0.9 mm). Functional and structural data were independently normalized into a common space (the Montreal Neurological Institute (MNI) template, IXI549Space) using SPM12 unified segmentation and normalization procedure with a reference functional image computed as the mean functional data after realignment across all timepoints omitting outlier scans. The output data were resampled to a common bounding box between MNI-space coordinates (−90, −126 and −72) and (90, 90 and 108), using 2 mm isotropic voxels and fourth-order spline interpolation for the functional data and 1-mm isotropic voxels and tri-linear interpolation for the structural data. Lastly, the functional data were smoothed spatially using spatial convolution with a 4 mm full-width half-maximum (FWHM) Gaussian kernel. For the language localizer task and the non-linguistic tasks, effects were estimated using a general linear model (GLM) in which each experimental condition was modeled with a boxcar function convolved with the canonical hemodynamic response function (HRF) (fixation was modeled implicitly). Temporal autocorrelations in the BOLD signal time series were accounted for by a combination of high-pass filtering with a 128 s cutoff and whitening using an AR(0.2) model (first-order autoregressive model linearized around the coefficient a = 0.2) to approximate the observed covariance of the functional data in the context of restricted maximum likelihood (ReML) estimation. In addition to main condition effects, other model parameters in the GLM design included first-order temporal derivatives for each condition (for modeling spatial variability in the HRF delays) as well as nuisance regressors to control for the effect of slow linear drifts, subject-motion parameters and potential outlier scans on the BOLD signal.

### fROI definition and response estimation

For each participant, language fROIs were defined using the Group-constrained Subject-Specific (GSS) approach (Fedorenko et al., 2010) whereby a set of parcels or ‘search spaces’ (that is, brain areas within which most individuals in prior studies showed activity for the localizer contrast) is combined with each individual participant’s activation map for the same contrast. (As noted above, in the main analysis, we used the reading-based language localizer, but see **Supp. Figure 1** for evidence that the results are similar when using the listening-based language localizer.)

We used five parcels derived from a group-level representation of data for the Sentences > Nonwords contrast in 220 independent participants. These parcels (used in much prior work; e.g., Malik-Moraleda, Ayyash et al., 2022) included three regions in the left frontal cortex—two in the inferior frontal gyrus (LIFG and LIFGorb) and one in the middle frontal gyrus (LMFG)— and two regions in the left lateral temporal cortex (LAntTemp and LPostTemp; **Figure 2**).

Individual fROIs were defined by selecting within each parcel the 10% of most localizer-responsive voxels based on the *t-*values for the relevant contrast (Sentences > Control). In our set of participants, 2 participants showed right-hemisphere dominance for language based on the English reading localizer. For these participants, we used the RH parcels to define the language fROIs, but we also report a version of the analysis where the LH parcels are used for all participants (**Supp. Figure 5**). We then extracted the responses from these fROIs (averaging the responses across the voxels in each fROI) to each condition of the language localizer (using an across-runs cross-validation procedure to ensure independence between the data used to define the fROIs vs. to examine the response magnitudes), to the Sentences and Control conditions of the auditory language localizer and of the critical conlang experiment, and to the non-linguistic tasks. Statistical tests were then performed on the percent BOLD signal change values extracted from the fROIs.

### Statistical analysis

For each of the tasks (and each conlang separately in some analyses), we fitted a model that predicted the BOLD responses (to the relevant contrasts, as elaborated below) of the LH language fROIs, with participants and fROIs included as random intercepts:

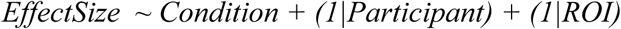

Note that for the main analyses, we consider the five language regions as an integrated system given that these regions a) have similar functional profiles with respect to their selectivity for language (e.g., Fedorenko et al., 2011; Fedorenko & Blank, 2020) and their role in lexico-semantic and combinatorial processing during language comprehension and production (e.g., Fedorenko et al., 2010, 2016, 2020; Bautista & Wilson, 2016; Hu, Small et al., 2022) and b) exhibit strong inter-region correlations in their activity during naturalistic cognition paradigms (e.g., Blank et al., 2014; Paunov et al., 2019; Braga et al., 2020; Malik-Moraleda, Ayyash et al., 2022).

To assess the selectivity of the language regions’ responses to language relative to control conditions, we examined the Sentences vs. Control contrast for the reading-based localizer (estimated using the across-runs cross-validation to ensure independence between the data used for fROI definition vs. response estimation), the listening-based localizer, and the critical conlang task both across all participants and within each conlang group. To assess the specificity of the language regions’ responses to conlangs relative to non-linguistic tasks, we examined the contrasts between the Sentences condition of the conlang task and each of the following conditions: the hard and easy conditions of the spatial working memory task and the mental and physical conditions from the naturalistic Theory of Mind task.

## Acknowledgements

We would like to thank Bozena Pajak, Jessica Becker, and Joseph Rollinson from Duolingo for providing the initial recordings for Esperanto, Klingon, and High Valyrian; Matthew Michael Early and Alan Anderson for re-recording the Esperanto and Klingon materials, respectively; So Hee Ahn and Rucha Kelkar for help with stimulus creation; and all members of the Fedorenko and Gibson labs for helping during data collection (especially, Colton Casto, Lia Washington, Aalok Sathe, Anya Ivanova, Hee So Kim, Eric Martinez, Sammy Floyd, Chengxu Zhuang, Moshe Poliak, Greta Tuckute, Hope Kean, Carina Kauf and Elizabeth Lee). We would also like to thank Damián Blasi and Arika Okrent for their presentations (along with the four conlang creators, who are authors on the manuscript) at the ‘Brains on Conlangs’ event at MIT (https://mcgovern.mit.edu/2022/12/12/brains-on-conlangs/), and Thomas Clark for helping host the event. Finally, we are grateful to Elise Malvicini, Julie Pryor, and Gayle Lutchen for their help in planning and organizing the event and to the McGovern Institute for Brain Research for sponsorship. For comments on an earlier draft, we thank Rebecca Saxe and Steve Piantadosi. This work was supported by research funds to EF from the McGovern Institute for Brain Research, the Brain and Cognitive Sciences Department, the Simons Center for the Social Brain, and the Middleton Professorship. EF was additionally supported by NIH awards R01-DC016607, R01-DC016950, and U01-NS121471.

