## Supplementary Information for "Constructed languages are processed by the same brain mechanisms as natural languages"

**Supplementary Text 1**: Procedure for the creation of acoustically degraded materials.

**Supplementary Table 1**: Demographic details and language background of the participants.

**Supplementary Table 2**: Phrases and sentences used in the critical conlang task and their English translations.

**Supplementary Table 3**: Details of the critical conlang experimental paradigm.

**Supplementary Table 4**: Common European Framework of Reference self-assessment grid (Council of Europe, 2001).

**Supplementary Table 5**: Results of the linear models fit separately for each of the conlang groups (i.e., Esperanto, Klingon, Na’vi, High Valyrian, Dothraki) examining the responses of the language areas to the Sentences and Control conditions of the language localizers and the critical conlang task.

**Supplementary Table 6**: Results of the linear models fit separately for each of the conlang groups (i.e., Esperanto, Klingon, Na’vi, High Valyrian, Dothraki) examining the responses of the language areas to the Sentences conditions of the conlang task and each of the four non-linguistic conditions.

**Supplementary Figure 1:** The language areas (defined by the Sentence>Control contrast in the native language listening task; cf. English) show a strong response during the processing of both natural and constructed languages.

**Supplementary Figure 2:** Each of the fROIs in the language network shows a strong response during the processing of both natural and constructed languages.

**Supplementary Figure 3:** The responses to Esperanto and four fictional conlangs in language regions are comparable.

**Supplementary Figure 4:** The amount of overlap (Dice coefficient) is robust to the percentage of voxels taken to define language-responsive voxels.

**Supplementary Figure 5:** The language areas (in the left hemisphere; cf. the language-dominant hemisphere) show a strong response during the processing of both natural and constructed languages.

**Supplementary Figure 6:** Whole-brain group-level activation maps for the non-linguistic tasks and responses to non-linguistic tasks in the Multiple Demand network and the Theory of Mind fROIs.

**Supplementary Figure 7:** Whole-brain individual-level activation overlap between conlang responses and responses to the language localizer vs. non-linguistic tasks.

**Supplementary Text 1: Procedure for the creation of acoustically degraded materials.**

To create the acoustically degraded versions of the linguistic recordings for use in the control conditions (for both the auditory version of the language localizer and the critical conlang experiments), we followed the procedure that was originally developed in Scott et al. (2017). In particular, the intact files were low-pass filtered at a pass-band frequency of 500 Hz. In addition, a noise track was created from each intact clip by randomizing 0.02 s long periods. To produce variation in the volume of the noise, the noise track was multiplied by the amplitude of the intact clip’s signal over time. The noise track was then low-pass filtered at a pass-band frequency of 8,000 Hz and a stop frequency of 10,000 Hz to soften the highest frequencies. The noise track and the low-pass-filtered copies of the intact files were then combined, and the level of noise was adjusted so as to render the clips unintelligible.

| **UID** | **Gender** | **Age** | **Hand** | **Proficiency** | **Num**  **Langs** | **Languages** |
| --- | --- | --- | --- | --- | --- | --- |
| *931* | M | 34 | R | 4 | 14 | English (0,20), F (23,17), M (14,16), **Esperanto (22,16),** K (19,14), J (16,10), P (27,9), S (5,9), T (25,8), C (16,7), G (31,6), A (22,5), G (14,4), P (26,4) |
| *938* | M | 37 | R | 5 | 3 | English (0,20), **Esperanto (30,16),** S (10,8) |
| *942* | M | 51 | R | 3 | 4 | English (0,20), **Klingon (27,12),** S (14,10), H (8,4) |
| *948* | M | 29 | R | 4 | 13 | English (0,20), **Esperanto (16,17),** F (17,14), D (0,12), L (6,11), G (6,10), A (4,10), ASL (4,10), S (6,9), I (16,9), T (20,7), A (6,5), M (14,4) |
| *949* | M | 33 | R | 4 | 4 | English (0,20), **Na’vi (27,16)**, J (9,9), R (26,4) |
| *950* | F | 50 | R | 2 | 3 | English (0,20), J (25,11), **High Valyrian (50,7)** |
| *952* | F | 31 | R | 3 | 4 | English (0,20), **High Valyrian (31,10),** S (12,10), H (5,8) |
| *953* | M | 18 | R | 5 | 6 | English (0,20), T (15,20), **Esperanto (14,19),** S (11,19), B (0,12), A (16,7) |
| *959* | M | 25 | R | 5 | 2 | English (0,20), **Esperanto (22,18)** |
| *960* | M | 18 | R | 5 | 4 | English (0,20), **Esperanto (18,17),** A (15,8), Y (19,8) |
| *961* | M | 21 | R | 4 | 4 | English (0,20), **Na’vi (15,15),** S (14,9), G (17,4) |
| *962* | M | 37 | R | 5 | 14 | English (5,20), S (0,19), **Esperanto (31,18),** F (32,14), I (13,11), P (33,10), E (35,9), H (33,8), G (35,8), J (30,7), R (35,6), S (33,6), P (32,6) |
| *963* | M | 27 | R | 4 | 4 | English (0,20), **Na’vi (22,16),** I (20,8), G (18,8) |
| *964* | M | 31 | R | 5 | 2 | English (0,20), **Na’vi (18,20)** |
| *965* | F | 25 | R | 5 | 3 | English (0,20), **Na’vi (20,16),** M (14,8) |
| *966* | F | 52 | R | NA | 10 | English (0,20), A (25,18), S (12,16), H (22,14), F (14,13), G (20, 13), P (19, 13), **Esperanto (34,11), Klingon** **(34, 9),** S (22,7) |
| *967* | F | 23 | R | 5 | 4 | English (0,20), **Esperanto (14,16),** S (14,11), J (14,11) |
| *968* | M | 30 | R | 5 | 7 | English (0,20), **Esperanto (22,20),** S (14,11), G (4,8), A (20,6), H (18,5) |
| *969* | M | 35 | R | 4 | 6 | English (0,20), G (10,15), **Na’vi (22,15),** M (28,10), S (8,9), T (20,4) |
| *970* | M | 39 | R | 5 | 2 | English (0,20), **Dothraki (32,15)** |
| *971* | F | 53 | R | 4 | 7 | English (0,20), **Klingon (48,14),** F (1,12), J (16,12), G (47,8), S (51,8), A (1,5) |
| *972* | M | 41 | R | NA | 25 | English (0,20), S (0,17), F (19,13), T (34, 16), A (14,7), **High Valyrian (32, 15)**, G (17,12), **Dothraki (28,17),** A (18, 10), E (18,14), R (18,8), F (34, 8), C (30, 13), H (19, 8), E (20,6), T (19,7), H (31,6), A (31,5), J (20,7), I (30,12), S (33,13), K (33,12), T (24,8), S (19,6), A (1,4) |
| *973* | M | 48 | R | 5 | 13 | English (0,20), **Esperanto (15,20),** S (6,15), P (37,12), F (7,12), G (5,10), R (20,10), L (12,9), W (45,8), G (44,8), T (37,8), B (45,8), G (13,7) |
| *974* | F | 23 | L | 4 | 4 | English (0,20), S (13,15), **Esperanto (19,14),** P (19,11) |
| *975* | F | 53 | R | 5 | 8 | English (0,20), **Klingon (23,20),** I (34,12), S (2,9), M (40,7), G (51,6), W (51,5), L (14,5) |
| *976* | F | 38 | R | NA | 6 | English (0,20), S (0,18), **Esperanto (28,14),** F (3,10), J (15,7), P (5,6) |
| *977* | M | 26 | L | 5 | 5 | English (0,20), **Esperanto (23,18),** M (25,7), I (25,7), S (14,6) |
| *978* | M | 42 | L | 4 | 5 | English (0,20), S (0,16), **Klingon (12,13),** F (0,7), M (16,5) |
| *979* | M | 26 | R | 4 | 2 | English (0,20), **Esperanto (17,12)** |
| *980* | M | 38 | R | 5 (Klingon) 3 (Esperanto) | 4 | English (0,20), **Klingon (7,17),** F (14,12), **Esperanto (15,12)** |
| *981* | M | 33 | R | 4 | 3 | English (0,20), **Klingon (15,14),** F (15,8) |
| *982* | M | 26 | L | 5 (Klingon)  3 (Esperanto) | 4 | English (0,20), F (12,20), **Klingon (21,20),** **Esperanto (22,15)** |
| *983* | F | 24 | R | 5 | 2 | English (0,20), **Esperanto (18,19)** |
| *984* | F | 31 | R | 4 | 5 | English (0,20), F (10,15), **Klingon (24,13),** S (10,9), N (30,5) |
| *985* | F | 29 | L | 4 | 3 | English (0,20), **Esperanto (24,18), S (3,13)** |
| *994* | F | 29 | R | 5 (Na’vi)  2 (Dothraki) | 11 | German (0,20), E (10,18), **Na’vi (18,14),** D (20,13), I (19,9), F (9,7), M (17,6), **Dothraki (20,6),** Q (22,4), S (22,4), B (22,2) |
| *995* | M | 31 | R | 4 | 11 | Dutch (0,20), E (8,19), **Na’vi (18,15),** G (12,14), F (12,9), J (20,9), L (12,5), A (12,5), I (18,5), L (17,4) |
| *996* | M | 24 | R | NA | 21 | English (0,20), S (20,20), S (9,18), F (16,18), W (22,15), I (20,15), G (19,14), I (16,13), P (NA,10), J (13,9), D (NA,9), A (14,8), A (27,8), G (16,8), H (27,8), T (30,8), C (NA,6), L (12,6)**, Klingon** (18,6), S (18,6), M (NA,5) |

**Supplementary Table 1**: **Demographic details and language background of the participants**. UID refers to the Unique ID assigned to the participant in the lab’s database (and can be cross-referenced with the data available at OSF: <https://osf.io/pe74v/>). Age refers to the participant’s age (in full years) at the time of testing. Hand refers to the participant’s handedness (R=right-handed; L=left- handed; A=ambidextrous). Proficiency refers to the proficiency for the relevant conlang(s) that participants reported during the initial contact (on an informal scale from 1 (“know a couple words”) to 5 (“can understand almost everything and express myself freely”). NumLangs refers to the number of languages that participants listed as having some proficiency in. Languages provides the list of languages, ordered by proficiency. The native language and the conlang(s) are listed by their full names (the conlang(s) is/are bolded); the remaining languages are abbreviated to the first letter in order to protect the participants’ identities. For each language, the first number in parentheses corresponds to the age of acquisition and the second number corresponds to overall proficiency on a scale from 0 to 20 (participants were asked to rate their ability in auditory comprehension, written comprehension, speaking, and writing on a scale from 0 (no knowledge) to 5 (native or native-like proficiency); the scores were summed to derive the overall score). Missing values are marked as NAs. A csv version of the table is available at OSF (<https://osf.io/pe74v/>).

| **Language** | **Item** | **Phrase/Sentence** | **English Translation** |
| --- | --- | --- | --- |
| Esperanto | 1 | Kiajn filmojn vi preferas? | What kinds of films do you prefer? |
| Esperanto | 2 | Ĉu mi renkontu vin ekster la biblioteko ? | Should I meet you outside the library? |
| Esperanto | 3 | Ĉu vi estas la gepatroj de Sofia? | Are you the parents of Sofia? |
| Esperanto | 4 | Kie vi aĉetis tiun manĝilaron? | Where did you buy that silverware? |
| Esperanto | 5 | Oni konstruis longan ponton super la larĝa rivero. | They built a long bridge over the wide river. |
| Esperanto | 6 | Fortranĉu pecon por mi! | Cut off a piece for me! |
| Esperanto | 7 | Ni povas influi nian socion per niaj agoj. | We can influence our society by our actions. |
| Esperanto | 8 | Vi bone progresas! | You are progressing well! |
| Esperanto | 9 | Ĝi estas tiel malgranda kiel formiko. | It is as small as an ant. |
| Esperanto | 10 | Indas eklabori nun. | It is worth it to start working now. |
| Esperanto | 11 | Mi sopiras al vi ĉiuj, inkluzive de la hundo! | I long for you all, including the dog! |
| Esperanto | 12 | Mi ne ŝatas labori nokte. | I don’t like to work at night. |
| Esperanto | 13 | Kiu estas la numero de tiu kanto? | Which is the number of that song? |
| Esperanto | 14 | Nia teamo havas ĉiajn fortojn. | Our team has all kinds of strengths. |
| Esperanto | 15 | Kiom da jaroj vi havas? | How old are you? |
| Esperanto | 16 | Ili havas belajn hundojn. | They have beautiful dogs. |
| Esperanto | 17 | Katido estas ido de kato. | A kitten is a cat’s offspring. |
| Esperanto | 18 | Ili malrapide kantas. | They sing slowly. |
| Esperanto | 19 | Mi parolas la hispanan nur iomete. | I only speak a little Spanish. |
| Esperanto | 20 | Mi ŝatas naĝadon. | I like swimming. |
| Esperanto | 21 | Bonvenon! | Welcome! |
| Esperanto | 22 | La domo de nia avo estas en Londono. | Our grandfather’s house is in London. |
| Esperanto | 23 | La flegisto havas grandan valizon. | The nurse has a large suitcase. |
| Esperanto | 24 | Kiam ŝi manĝas oranĝojn? | When does she eat oranges? |
| Esperanto | 25 | Kiom da greno estas en os hi? | How much grain is in the container? |
| Esperanto | 26 | Ĉu vi volas fariĝi la urbestro de via urbo? | Do you want to become the mayor of your city? |
| Esperanto | 27 | Vi devus malfermi vian menson. | You should open your mind. |
| Esperanto | 28 | Pri kio la ĉevalo demandas? | What is the horse asking about? |
| Esperanto | 29 | Mi pensas, ke ne estos eraro. | I think there will be no mistake. |
| Esperanto | 30 | Ĉu vi povos viziti nin en oktobro? | Will you be able to visit us in October? |
| Esperanto | 31 | Ĉu vi freŝe kuiris la matenmanĝon ? | Have you just cooked the breakfast? |
| Esperanto | 32 | Ni laboras kune en spirito de frateco. | We work together in a spirit of brotherhood. |
| Esperanto | 33 | Kantu al ni en Esperanto ! | Sing to us in Esperanto! |
| Esperanto | 34 | La tondro timigis min. | The thunder scared me. |
| Esperanto | 35 | Ni ŝatas fromaĝon kun vino. | We like cheese with wine. |
| Esperanto | 36 | Naŭdek estas pli ol kvardek. | Ninety is more than forty. |
| Esperanto | 37 | La instruisto havas diversajn ideojn. | The teacher has diverse ideas. |
| Esperanto | 38 | Post la kurso, la partoprenintoj iris al kafejo kune. | After the course, the participants went to a cafe together. |
| Esperanto | 39 | Ĉu vi amas ŝin aŭ min? | Do you love her or me? |
| Esperanto | 40 | Miaj gepatroj volas, ke ni prokrastu la vespermanĝon, ĉar neĝas. | My parents want us to postpone the dinner because it is snowing. |
| Esperanto | 41 | Ili ofte parolas pri scienco dum siaj podkastoj. | They often speak about science during their podcasts. |
| Esperanto | 42 | Ĝirafoj kutime tute ne bojas. | Giraffes usually do not bark at all. |
| Esperanto | 43 | Lidja Zamenhof lernis Esperanton kiel naŭ-jara knabino. | Lidja Zamenhof learned Esperanto as a nine-year-old girl. |
| Esperanto | 44 | Estas nekredeble! | It is incredible! |
| Esperanto | 45 | Mia mantelo estas tro larĝa. | My coat is too wide. |
| Esperanto | 46 | Li iris al Londono kun sia kunulo. | He went to London with his partner. |
| Esperanto | 47 | Jen hundo. Ĝia nomo estas Sofia. | Here is a dog. Its name is Sofia. |
| Esperanto | 48 | Oni pensas per sia cerbo. | One thinks with one’s brain. |
| Esperanto | 49 | Post tiu kiso, ĉu vi amas min? | After that kiss, do you love me? |
| Esperanto | 50 | Ĉu tiu planto estas manĝebla? | Is that plant edible? |
| Esperanto | 51 | Tio estas mia fina decido. | That is my final decision. |
| Esperanto | 52 | Multaj lingvoj diferencas de la nia. | Many languages differ from ours. |
| Esperanto | 53 | Li flaras la vespermanĝon. | He smells the evening meal. |
| Esperanto | 54 | Mi estas nenies malamiko. | I am nobody’s enemy. |
| Esperanto | 55 | Ju malpli da vortoj, des pli bone. | The fewer words, the better. |
| Esperanto | 56 | Ĉu vi estas Sinjoro Mango? | Are you Mr. Mango? |
| Esperanto | 57 | Frago estas frukto. | A strawberry is a fruit. |
| Esperanto | 58 | La najbaro jam sidas sur la sofo. | The neighbor is already sitting on the sofa. |
| Esperanto | 59 | Ĉu tiu avino vere disdonis la sablerojn? | Did that grandmother really distribute the grains of sand? |
| Esperanto | 60 | Ne enuigu nin denove per tiu rakonto! | Don’t bore us again with that story! |
| Esperanto | 61 | La vespermanĝo estas fiŝaĵo kaj supo. | The dinner is fish and soup. |
| Esperanto | 62 | Elsalutu kaj iru labori! | Log off and go to work! |
| Esperanto | 63 | Ĉu ni estas bonaj patroj ? | Are we good fathers? |
| Esperanto | 64 | Kial vi ĉiam ridas? | Why are you always laughing? |
| Esperanto | 65 | Mateno estas antaŭ tagmezo. | Morning is before noon. |
| Esperanto | 66 | Tero moviĝas ĉirkaŭ la suno. | Earth moves around the sun. |
| Esperanto | 67 | Li havas bonan apetiton. | He has a good appetite. |
| Esperanto | 68 | Al Toronto flugas multaj aviadiloj. | Many planes fly to Toronto. |
| Esperanto | 69 | Nepre ne kredu tion! | Absolutely dont believe that! |
| Esperanto | 70 | Vi povas lerni Esperanton en la reto. | You can learn Esperanto on the net. |
| Esperanto | 71 | Vi surhavas miajn botojn. | You are wearing my boots. |
| Esperanto | 72 | La enirejo situas dek metrojn post tiu strato. | The entrance is located ten meters after that street. |
| Esperanto | 73 | Ĉu estos frosto morgaŭ? | Will there be frost tomorrow? |
| Esperanto | 74 | Ŝi iris al Novjorko por kunveno. | She went to New York for a meeting. |
| Esperanto | 75 | Saluton, Sofia! | Hello, Sofia! |
| Esperanto | 76 | Kio estas la simbolo de tiu partio? | What is the symbol of that party? |
| Esperanto | 77 | Niaj du hundoj kaj tri katoj estas en la domo. | Our two dogs and three cats are in the house. |
| Esperanto | 78 | La knabino kantis elstare. | The girl sang outstandingly. |
| Esperanto | 79 | Estas pli ol mil denaskaj parolantoj de Esperanto. | There are more than a thousand native speakers of Esperanto. |
| Esperanto | 80 | Kiom da lano mi devas aĉeti por triki ĉapelon? | How much yarn do I need to buy to knit a hat? |
| Esperanto | 81 | Ni nomiĝas la verdaj steloj. | We call ourselves the green stars. |
| Esperanto | 82 | La flago de lia infano estas blua kaj ruĝa. | His child’s flag is blue and red. |
| Esperanto | 83 | Kvankam ŝi ne parolas la japanan, ŝi tamen parolas la ĉinan. | Although she does not speak Japanese, she does speak Chinese. |
| Esperanto | 84 | Malŝaltu la lampon ! Mi volas dormi. | Turn off the lamp! I want to sleep. |
| Esperanto | 85 | Ĉu vi havas liberan tempon? | Do you have free time? |
| Esperanto | 86 | Ŝi kapablas flue paroli ses lingvojn. | She can speak six languages fluently. |
| Esperanto | 87 | Kie oni povas aĉeti ian nigran robon ? | Where can one buy some kind of black dress? |
| Esperanto | 88 | La bestoj eniris la grandan boaton duope, ne unuope. | The animals entered the big boat two at a time, not one at a time. |
| Esperanto | 89 | Tridek minus dek estas dudek. | Thirty minus ten is twenty. |
| Esperanto | 90 | La tri lacaj beboj dormas. | The three tired babies are sleeping. |
| Esperanto | 91 | Estas tro malfrue. | It’s too late. |
| Esperanto | 92 | La sableroj en mia ŝuo ĝenas min. | The grains of sand in my shoe annoy me. |
| Esperanto | 93 | Multaj klubanoj manĝis kune en nia hejmo. | Many club members ate together in our home. |
| Esperanto | 94 | Nia domo enhavas kvar ĉambrojn. | Our house contains four rooms. |
| Esperanto | 95 | Ĉu io malbona okazis? | Did something bad happen? |
| Esperanto | 96 | La loĝantaro de Usono estas pli ol tricent milionoj da homoj. | The population of the United States is more than three hundred million people. |
| Esperanto | 97 | Adamo estas lia nomo. | Adam is his name. |
| Esperanto | 98 | Ĉu la glaso estas plena aŭ malplena? | Is the glass full or empty? |
| Esperanto | 99 | La akvo fluis de la monto al la maro. | The water flowed from the mountain to the sea. |
| Esperanto | 100 | La knabo estas malkontenta, ĉar li ne havas katon. | The boy is discontented because he does not have a cat. |
| Klingon | 1 | torgh, tlhIngan yaj’a’ mara? | Torg, did Mara understand the Klingon? |
| Klingon | 2 | toDujlIjmo’ qavuv. | Because of your courage, I respect you. |
| Klingon | 3 | SoSnI’pu’lI’ | your grandmothers |
| Klingon | 4 | wo’rIv luvoq torgh mara je. | Torg and Mara trusted Worf. |
| Klingon | 5 | HibIjQo’! | Don’t punish me! |
| Klingon | 6 | nuqDaq ‘oHtaH baghneQ’e’? | Where is the spoon? |
| Klingon | 7 | mughIj HoSDo’. | Energy beings scare me. |
| Klingon | 8 | targhmey bIH. | They are targs. |
| Klingon | 9 | SoSnI’, qatlh Ho’Du’ law’ Daghaj? | Grandmother, why do you have many teeth? |
| Klingon | 10 | Haqtajvam boch law’ jul boch puS. | This scalpel is shinier than the sun. |
| Klingon | 11 | paq wa’maHDIch yIlaD! | Read the tenth book! |
| Klingon | 12 | wa’Hu’ runpI’ chu’ vIje’. | I bought a new teapot yesterday. |
| Klingon | 13 | reH wo’vaD Suvqang tlhIngan SuvwI’. | A Klingon warrior is always willing to fight for the empire. |
| Klingon | 14 | SaH ghojmoHwI’. | The teacher is present. |
| Klingon | 15 | qet torgh mara je. | Torg and Mara run. |
| Klingon | 16 | HoDpu’ tISam! | Find the captains! |
| Klingon | 17 | muquvmoH puqnI’loDma’ puqnI’be’ma’ je. | Our grandson and our granddaughter honor me. |
| Klingon | 18 | cha’maH | twenty |
| Klingon | 19 | nov vuv qIrq HoD. | Captain Kirk respected the alien. |
| Klingon | 20 | puq SopmoH vav. | The father feeds the child. |
| Klingon | 21 | pa’ jaghpu’ law’ tu’lu’. | There are many enemies there. |
| Klingon | 22 | De’wI’lIjDaq Qorwaghmey tu’lu’’a’? ghobe’. Tera’ jentu’ tu’lu’. | Are there windows in your computer? No. There is a penguin. |
| Klingon | 23 | loS loS loD. | The four men waited. |
| Klingon | 24 | bIvItmo’ qabIjbe’. | Because you told the truth, I will not punish you. |
| Klingon | 25 | pe’vIl peSuv! | Fight forcefully! |
| Klingon | 26 | wa’Hu’ Dilegh. | We saw them yesterday. |
| Klingon | 27 | Qup ‘ej Qip. | He is young and stupid. |
| Klingon | 28 | tera’nganpu’vetlh | those Terrans |
| Klingon | 29 | wo’rIv ‘IH law’ torgh ‘IH puS. | Worf is more handsome than Torg. |
| Klingon | 30 | ‘arlogh Dalo’pu’? | How many times have you used it? |
| Klingon | 31 | taj ‘oH’a’ Dochvam’e’? | Is this a knife? |
| Klingon | 32 | bIQIp’a’? ghobe’. jIval. | Are you stupid? No. I am smart. |
| Klingon | 33 | tlhInganpu’ | the Klingons |
| Klingon | 34 | bImoH, ghotI’ Darur. | You are as ugly as a fish. |
| Klingon | 35 | SirlIy wISamlaHbe’. | We cannot find any silver. |
| Klingon | 36 | tlhInganpu’ chaH mara’e’ wo’rIv’e’ je. | Mara and Worf are Klingons. |
| Klingon | 37 | muDuQ ghuQvam. | This poem speaks to me. |
| Klingon | 38 | moH DaSmeylIj. moH je poghmeylIj. | Your boots are ugly. Your gloves are also ugly. |
| Klingon | 39 | Qivam luchenmoH qama’. | Prisoners built this bridge. |
| Klingon | 40 | tIn Duj ‘ej mach qach. | The ship is big and the building is small. |
| Klingon | 41 | yuQ ‘oH tera’’e’. | Earth is a planet. |
| Klingon | 42 | targh ghIj wamwI’ Soy’. | The clumsy hunter frightened the targ. |
| Klingon | 43 | ngop Hivje’mey je | plates and glasses |
| Klingon | 44 | barat Dab ‘IrneHwI’. | My maternal uncle lives in India. |
| Klingon | 45 | qagh tujmoH Human vutwI’. | The human chef heated the gagh (a delicacy of worms). |
| Klingon | 46 | tajwIj jejHa’moH ‘Iv? | Who made my knife blunt? |
| Klingon | 47 | Dulegh’a’ vay’? ghobe’. Mulegh pagh. | Did anybody see you? No. Nobody saw me. |
| Klingon | 48 | Sarghmey | sarks (a horse-like animal) |
| Klingon | 49 | SoHvaD Dochmeyvam nobbe’ loDpu’. | The men did not give you these things. |
| Klingon | 50 | jImach. | I am little. |
| Klingon | 51 | torgh ‘oH loD pong’e’. | The man’s name is Torg. |
| Klingon | 52 | qama’vaD baghneQvam yInob! | Give this spoon to the prisoner! |
| Klingon | 53 | yInSIp lupoQ Depmeyvam. | These beings require oxygen. |
| Klingon | 54 | qach ‘emDaq QamtaH lurveng ‘ej tlhIch purtaH. | Lurveng is standing behind the building and smoking. |
| Klingon | 55 | bejwI’pu’ bej ‘Iv? | Who watches the watchers? |
| Klingon | 56 | yIvbeH | a tunic |
| Klingon | 57 | ‘Iw Hiq botlhutlh tlhIH. | You drank the bloodwine. |
| Klingon | 58 | maH | we |
| Klingon | 59 | qagh jab jabwI’. | The waiter served gagh. |
| Klingon | 60 | wa’ tlhay | one sleeve |
| Klingon | 61 | ‘IQ verengan tlhIngan Hol jatlhlaHbe’mo’. | The Ferengi is sad because she cannot speak Klingon. |
| Klingon | 62 | jIyIn. | I live. |
| Klingon | 63 | Hut Qebmey ghaj jubbe’wI’pu’. | The mortals had nine rings. |
| Klingon | 64 | Sughungchugh vaj naHmey tISop! | If you are hungry then eat vegetables! |
| Klingon | 65 | qoq tIn tI’ qoq mach. | The small robot repaired the big robot. |
| Klingon | 66 | Say’’eghmoH yaS. | The officer cleaned himself. |
| Klingon | 67 | ghoqwI’ ghIj boQwI’ HoSghaj. | The powerful assistant scared the spy. |
| Klingon | 68 | rojmabmo’ reHoHbe’. | Because of the peace treaty, we won’t kill you. |
| Klingon | 69 | SubuDmo’ Suluj. | Because you are lazy, you failed. |
| Klingon | 70 | wej Qorwaghmey vISoQmoHtaH. | I am closing the three windows. |
| Klingon | 71 | ‘al’on yIghor! | Break the glass! |
| Klingon | 72 | tlhIngan jIH’a’? ghobe’. Tera’ngan SoH. | Am I a Klingon? No. You are a Terran. |
| Klingon | 73 | Qapla’ batlh je! | Success and honor! |
| Klingon | 74 | chu’ jengva’vam. | This plate is new. |
| Klingon | 75 | paqlIj | your book |
| Klingon | 76 | SaH ‘Iv? | Who cares? |
| Klingon | 77 | SopDI’ mara, wa’maH chorghlogh Qoylu’pu’. | Mara will eat at eighteen hundred hours. |
| Klingon | 78 | tlhImwIjDaq qungmey mach chenmoH ghewmey. | The bugs made little holes in my carpet. |
| Klingon | 79 | jIwuQchoHtaH. | I am getting a headache. |
| Klingon | 80 | baghneQmey chIS vIghaj. | I have the white spoons. |
| Klingon | 81 | tI’’egh verenganpu’ Duj. | The Ferengi’s ship repaired itself. |
| Klingon | 82 | targhmeyvam HoH ‘Iv? | Who killed these targs? |
| Klingon | 83 | yuDHa’ ghaH. | She is honest. |
| Klingon | 84 | jengva’vetlh | that plate |
| Klingon | 85 | rop loDnalwI’. | My husband is ill. |
| Klingon | 86 | malja’maj qangtlhIn qolqoS ‘oH yuDHa’ghach’e’. | Honesty is the most important part of our company’s philosophy. |
| Klingon | 87 | Hipup! | Kick me! |
| Klingon | 88 | Duy’’a’ qoqlIj? | Is your robot defective? |
| Klingon | 89 | tlhIngan SoH. | You are a Klingon. |
| Klingon | 90 | SoSwI’ ghaHbe’ vavwI’ be’nal’e’. | My father’s wife is not my mother. |
| Klingon | 91 | ngemDaq ghach Ha’DibaHmey Qob. | Dangerous animals lurk in the forest. |
| Klingon | 92 | qoqmey | robots |
| Klingon | 93 | wa’vatlh cha’maH wej | one hundred twenty-three |
| Klingon | 94 | vongwI’pu’ tIbIjQo’! | Don’t punish the hypnotists! |
| Klingon | 95 | qIbvam | this galaxy |
| Klingon | 96 | yIn ‘u’ Hoch je | life, the universe, and everything |
| Klingon | 97 | pa’ boqor HoD Dalegh’a’? | Did you see Captain Bokor there? |
| Klingon | 98 | yIn mara. | Mara lives. |
| Klingon | 99 | ngo’qu’ paqvetlh. | That book is very old. |
| Klingon | 100 | bItlhutlhnIS’a’? HISlaH. jItlhutlhnIS. | Do you need to drink? Yes. I need to drink. |
| Na’vi | 1 | Sunu oer fwa slele. | I like to swim. |
| Na’vi | 2 | Ayfo kempe si mì na’rìng ? | What do they do in the forest? |
| Na’vi | 3 | Omum oel teyngta pelun pol ke tok fìtsenget. | I know why she isn’t here. |
| Na’vi | 4 | Lu fìkilvan sloa nìhawng; ke tsun fko emkivä. | This river is too wide to cross. |
| Na’vi | 5 | Lu poe sevin nìftxan kuma yawne slolu oer. | She was so beautiful, I fell in love with her. |
| Na’vi | 6 | Sunu oeyä palukantsyìpur fwa yom payoangit. | My cat likes to eat fish. |
| Na’vi | 7 | Tsa’opin a sunu oer frato lu eampin. | My favorite color is blue. |
| Na’vi | 8 | Ke sunu oer fwa tìkangkem si txonkrr. | I don’t like to work at night. |
| Na’vi | 9 | Oeti nìn rä’ä tsafya! | Don’t look at me like that! |
| Na’vi | 10 | Lu foru aynantangtsyìp alor. | They have beautiful dogs. |
| Na’vi | 11 | Oeri solalew zìsìt apxevol. | I’m 24 years old. |
| Na’vi | 12 | Txo nga ‘efu ohakx, oeng yivom ko! | If you’re hungry, let’s eat! |
| Na’vi | 13 | Poe lu tsulfätu lì’fyayä leNa’vi. | She is a master of the Na’vi language. |
| Na’vi | 14 | Tsaolo’eyktan lu tutan asìltsan. | The clan leader is a good man. |
| Na’vi | 15 | Ayoe tìkangkem si ‘awsiteng nìsoaia. | We work together as a family. |
| Na’vi | 16 | Sìlpey oe tsnì fìtìrol sìyevunu ngar. | I hope you’ll like this song. |
| Na’vi | 17 | Lu oeru sngum a fo ke tìyevätxaw. | I’m worried that they won’t return. |
| Na’vi | 18 | Oel pukit fpole’ ngaru trram. | I sent you the book yesterday. |
| Na’vi | 19 | Oeyä ‘Eylan alu Sofia kelku si mì Nu Yorkì. | My friend Sofia lives in New York. |
| Na’vi | 20 | Aylì’fya eltur tìtxen si nìtxan. | Languages are very interesting. |
| Na’vi | 21 | Po tìkangkem ke si pelun? | Why isn’t he working? |
| Na’vi | 22 | Furia nì’Ìnglìsì pamrel si oe, oeru txoa livu. | I’m sorry for writing in English. |
| Na’vi | 23 | Hivahaw nìmwey, ma paskalin. | Sleep well, honey. |
| Na’vi | 24 | Polawm po san srake Neytiri holum sìk. | He asked if Neytiri left. |
| Na’vi | 25 | Oe new ngeyä tsmukehu muntxa sivi. | I want to marry your sister. |
| Na’vi | 26 | Fwa tok fìtsenget lu oeru meuia. | It’s an honor for me to be here. |
| Na’vi | 27 | Fìtìpe’un ke layatem. | This decision will not change. |
| Na’vi | 28 | Lu mì tampay payoang apxay. | There are many fish in the sea. |
| Na’vi | 29 | Peng oer teyngta lumpe nga sti. | Tell me why you’re angry. |
| Na’vi | 30 | ‘Ul yom, ‘ul slu txur. | The more you eat, the stronger you’ll get. |
| Na’vi | 31 | Tsatìoeyktìng srung soli oer fte tìngäzìkit tslivam. | That explanation helped me understand the problem. |
| Na’vi | 32 | Oe mowar si ngaru tsnì tsyivul pxiye’rìn. | I advise you to begin immediately. |
| Na’vi | 33 | Tsavur eltur tìtxen ke si. | That story isn’t interesting. |
| Na’vi | 34 | Lu fìkilvanmì spono alor. | There’s a beautiful island in this river. |
| Na’vi | 35 | Ke tsun fko tsawket tsive’a txonkrr. | One can’t see the sun at night. |
| Na’vi | 36 | ‘Rrta pxaw tsawke rikx. | The earth moves around the sun. |
| Na’vi | 37 | Oeri re’o tìsraw sängi. | My head hurts. |
| Na’vi | 38 | Nga krrpe hu karyu ultxa sayi? | When will you meet the teacher? |
| Na’vi | 39 | Rutxe foti rä’ä spivaw! | Please don’t believe them! |
| Na’vi | 40 | Sivop nìzawnong, ma ‘ite. | Have a good trip, daughter. |
| Na’vi | 41 | Oeru peyä tsmukan snumìna latsam. | His brother seems dim to me. |
| Na’vi | 42 | Tewti, nga kanu lu fìtxan nang! | Gee, you’re so smart! |
| Na’vi | 43 | Pol polawm tenga tìpawmit alo amrr. | She asked the same question five times. |
| Na’vi | 44 | Menantangtsyìp sì pxefalukantsyìp yerom. | Two dogs and three cats are eating. |
| Na’vi | 45 | Tsa’evenge tsun rivol nìmiklor. | That girl can sing beautifully. |
| Na’vi | 46 | Tsafkxilet tolìng ngaru tupel? | Who gave you that necklace? |
| Na’vi | 47 | Plltxe frapo san fìmauti lu ftxìlor sìk. | Everybody says this fruit is delicious. |
| Na’vi | 48 | Ftxozäri aylrrtok ngaru, ma sempu ! | Happy birthday, daddy! |
| Na’vi | 49 | Lu oeru fmawn a tsranten. | I have some important news. |
| Na’vi | 50 | Kxì, ma ‘eylan ! Kempe leren ? | Hey, dude! What’s up? |
| Na’vi | 51 | Po plltxe nìNa’vi nìwin na hufwe. | He speaks Na’vi fluently. |
| Na’vi | 52 | Lumpe poltxe nga san ke sunu oer fwa taron sìk? | Why did you say you don’t like to hunt? |
| Na’vi | 53 | Ke lu kawtu ro helku. | No one is at home. |
| Na’vi | 54 | Ayyayo rerol. | The birds are singing. |
| Na’vi | 55 | Lu tsa’evenganur ‘eylan. | The boy has a friend. |
| Na’vi | 56 | Ayfrrtu herahaw tsatseng. | The guests are sleeping there. |
| Na’vi | 57 | Tsa’evengel tìng syulangit taronyur. | The girl is giving flowers to the hunter. |
| Na’vi | 58 | Tsatutan lu sempul oeyä. | That man is my father. |
| Na’vi | 59 | Tsatxanlokxemì, aysutan sì aysuté ke ftia ‘awsiteng. | In that country, men and women do not study together. |
| Na’vi | 60 | Ke omum teyngta Ralu kolä pesengne. | I don’t know where Ralu went. |
| Na’vi | 61 | Lu ngeyä sempulur mepun atsawl. | Your father has big arms. |
| Na’vi | 62 | Sneyä sa’nok yawne lu tsa’evenganur. | The boy loves his mother. |
| Na’vi | 63 | Fì’evengan ke lu ‘itan oeyä. | This boy is not my son. |
| Na’vi | 64 | Pefnerel arusikx sunu ngar frato? | What kinds of films do you prefer? |
| Na’vi | 65 | Nga ngong längu, ha ke flolä. | You’re lazy, so you didn’t succeed. |
| Na’vi | 66 | Srake payoang tsun yivom yayot? | Can a fish eat a bird? |
| Na’vi | 67 | Zamunge oel ‘upxaret a ta Eywa. | I bring a message from Eywa. |
| Na’vi | 68 | Ngal pelun ngeyä tsko swizawti tolìng oeru? | Why did you give me your bow and arrow? |
| Na’vi | 69 | Kelku feyä hì’i lu nìtxan. | Their home is very small. |
| Na’vi | 70 | Ngeyä tìpawmìri tolel ngal tì’eyngit srak? | Did you receive an answer to your question? |
| Na’vi | 71 | Fyape fko syaw ngar, ma ‘evi? | What’s your name, little girl? |
| Na’vi | 72 | Fì’eveng ahì’i ke tsun ikranit mivakto. | This little child cannot ride a banshee. |
| Na’vi | 73 | Prrnen ke tsun pivlltxe. | A baby cannot speak. |
| Na’vi | 74 | Ngeyä tsmukanìl tok pesenget set? | Where is your brother now? |
| Na’vi | 75 | Ngal pelun pot spolaw? | Why did you believe him? |
| Na’vi | 76 | Fìlì’fyavi lesar lu nìtxan. | This expression is very useful. |
| Na’vi | 77 | Rutxe palukantsyìpur yomtìng trray. | Please feed the cat tomorrow. |
| Na’vi | 78 | Tsa’evengìl ahì’i ke tslam fya’ot a inan pamrelti. | The little child doesn’t know how to read. |
| Na’vi | 79 | Nga lu le’awa tute a tsun srung sivi. | You are the only one who can help. |
| Na’vi | 80 | Sko frrtu a mì helku ngeyä, oe ‘efu nitram frakrr. | I always feel happy as a guest in your home. |
| Na’vi | 81 | Ngaru lu fpom fìtrr srak? | How are you today? |
| Na’vi | 82 | Srake syuve som lu nìhawng? | Is the food too hot? |
| Na’vi | 83 | Feyä lì’fyati oel ke tslam. | I don’t understand their language. |
| Na’vi | 84 | Peyä sa’nok ayawne tolerkängup trram. | His dear mother died yesterday. |
| Na’vi | 85 | Faylì’uru tìng mikyun. | Listen to these words. |
| Na’vi | 86 | Talun tìtstew ngeyä, oe ngaru leioae si. | Because of your courage, I respect you. |
| Na’vi | 87 | Tsaspe’etut lonu pxiset! | Release the captive now! |
| Na’vi | 88 | Oeyä ‘eylan lu spxin nìtxan. | My friend is very sick. |
| Na’vi | 89 | Awnga ne na’rìng kivä ko fte stivarsìm aysyulangit. | Let’s go to the forest to gather flowers. |
| Na’vi | 90 | Fìstxeliri akosman irayo. | Thank you for this wonderful gift. |
| Na’vi | 91 | Ayoel fot tsole’a trram. | We saw them yesterday. |
| Na’vi | 92 | Nga zene fnivu set. | You have to be quiet now. |
| Na’vi | 93 | Tivel ngal ta prrnen ngeyä lawnolti. | May your little one bring you great joy. |
| Na’vi | 94 | Fìtaronyu lu txur sì tstew. | This hunter is strong and brave. |
| Na’vi | 95 | Tsa’u lu tsole’a oel a utral a tsawl frato mì sìrey. | That’s the tallest tree I’ve ever seen in my life. |
| Na’vi | 96 | Txopu rä’ä si, tsa’u lu tsawla palukantsyìp nì’aw. | Don’t be afraid, it’s just a big cat. |
| Na’vi | 97 | Nì’i’a leiu tsam hasey. | The war is finally over. |
| Na’vi | 98 | Po ke new slivu tsamsiyu. | He doesn’t want to become a warrior. |
| Na’vi | 99 | Lam oer fwa tsatutan ‘efu keftxo nìtxan. | That man seems very upset. |
| Na’vi | 100 | Krro len ayhem afe’. | Sometimes bad things happen. |
| HighValyrian | 1 | Ābra ūī ñuho velmot irughas. | The woman is giving it to my aunt. |
| HighValyrian | 2 | Daorys kesīr ilza. | There is no one here. |
| HighValyrian | 3 | Naejon eman. | I have a torso. |
| HighValyrian | 4 | Sparos Īliliot Zenturliot ilza? | Who is at the Inn at the Crossroads? |
| HighValyrian | 5 | Sēter rȳbon daor. | I do not hear the spell. |
| HighValyrian | 6 | Sylvie vala kostōbī ābrī majaqsa. | The wise man admires powerful women. |
| HighValyrian | 7 | Ampa taobi sīkudī ampā riñī jorrāelzi. | Ten boys love seventeen girls. |
| HighValyrian | 8 | Averilloma voktys daoriot istas. | The drunk priest went nowhere. |
| HighValyrian | 9 | Zenturliot hen kisalbrot umban. | I am staying at the inn for the feast. |
| HighValyrian | 10 | Issa, kepa iksan. | Yes, I am a father. |
| HighValyrian | 11 | Skoriot bona taoba vāedas? | Where is that boy singing? |
| HighValyrian | 12 | Ñuhi yrgos pamā? | Are you rubbing my neck? |
| HighValyrian | 13 | Taobi zoklī jorrāelzi. | The boys love the wolves. |
| HighValyrian | 14 | Kepa riñe rijas. | The father praises the girl. |
| HighValyrian | 15 | Taoba riñe urnes. | The boy sees a girl. |
| HighValyrian | 16 | Melo prūbroti iā kasto prūbroti vaoresā? | Do you prefer red apples or green apples? |
| HighValyrian | 17 | Taoba zȳhi gevie yrgos ūndas. | The boy saw his beautiful neck. |
| HighValyrian | 18 | Ñuha muña Dovaogēdī rijas. | My mother is praising the Unsullied. |
| HighValyrian | 19 | Hontesse vāedis! | The birds are singing! |
| HighValyrian | 20 | Zaldrīzesse hae āeksiot yne iotāptīlzi. | The dragons will consider me to be the master. |
| HighValyrian | 21 | Kesi kirini korzi issi? | Are these happy swords? |
| HighValyrian | 22 | Taoba raqiros ēza. | The boy has a friend. |
| HighValyrian | 23 | Sparos Taena issa? | Who is Taena? |
| HighValyrian | 24 | Ñuhi hāri velmanni bȳriar yba ēzi. | My three cousins have six knees. |
| HighValyrian | 25 | Ñuhi anni kostōbī zoklī urnesi. | My horses see the powerful wolves. |
| HighValyrian | 26 | Zentyssy konīr ēdrusi. | The guests are sleeping there. |
| HighValyrian | 27 | Dāria Thorot bardutas. | The queen wrote to Thoros. |
| HighValyrian | 28 | Jaehoti azantī izūgan. | I fear the knights of the gods. |
| HighValyrian | 29 | Ñuhi muñi nagesi. | My mothers are sweating. |
| HighValyrian | 30 | Ñuhe geltī pilvō daor? | Are you not holding my helmet? |
| HighValyrian | 31 | Riña arghurot rūkla irughas. | The girl is giving flowers to the hunter. |
| HighValyrian | 32 | Bona vala ñuha kepa issa. | That man is my father. |
| HighValyrian | 33 | Tyrion Iēmio dubys issa. | Tyrion is Jaime’s sibling. |
| HighValyrian | 34 | Vali ābrommi iprattis. | The men ate with the women. |
| HighValyrian | 35 | Aōha qȳbranna rōvi qimos ēza. | Your cousin has a large chin. |
| HighValyrian | 36 | Kesi gerpi issi. | These ones are the fruits. |
| HighValyrian | 37 | Taoba dōre vokti majaqsa. | The boy admires no priest. |
| HighValyrian | 38 | Dovaogēdī rȳban? | Do I hear the Unsullied? |
| HighValyrian | 39 | Dārilaros avera qurdot nevetas. | The prince carried the grapes to the table. |
| HighValyrian | 40 | Atroksia ipradas. | The owl eats. |
| HighValyrian | 41 | Taoba muñe urnes. | The boy sees the mother. |
| HighValyrian | 42 | Qumblie geltī emā. | You have a thick helmet. |
| HighValyrian | 43 | Taoba ñuha trēsy iksos daor. | The boy is not my son. |
| HighValyrian | 44 | Drīvose daor. | Not actually. |
| HighValyrian | 45 | Kesi ñurha sētera issi. | These ones are my magic spells. |
| HighValyrian | 46 | Vala iprattas se nagetas. | The man ate and sweated. |
| HighValyrian | 47 | Drīvose, ēdrī. | Actually, we are sleeping. |
| HighValyrian | 48 | Hontes klios ipradas. | The bird is eating the fish. |
| HighValyrian | 49 | Ñuhon lenton rōvon issa. | My house is big. |
| HighValyrian | 50 | Daenerys lōgra urnes. | Daenerys sees the ships. |
| HighValyrian | 51 | Embrī rȳbis. | They hear the oceans. |
| HighValyrian | 52 | Idañi issi. | They are twins. |
| HighValyrian | 53 | Konon lenton rōvon issa. | That house is large. |
| HighValyrian | 54 | Melisandre Iōnos majaqsa. | Melisandre admires John. |
| HighValyrian | 55 | Rȳbā? Daenerys ȳdras. | Do you hear? Daenerys is speaking. |
| HighValyrian | 56 | Vala atroksie rȳbas. | The man hears the owl. |
| HighValyrian | 57 | Riña Dovaogēdo muñe rhaenas. | The girl finds the Unsullied’s mother. |
| HighValyrian | 58 | Dārōñar qintra izūgan. | I fear the royal turtles. |
| HighValyrian | 59 | Mirros urnē? | Do you see something? |
| HighValyrian | 60 | Valyry korzī ēza! | The Valyrian has a sword! |
| HighValyrian | 61 | Āeksia qilōnommi pōnte idakosi! | The masters are attacking them with whips! |
| HighValyrian | 62 | Melva sinditon daor. | I did not buy pears. |
| HighValyrian | 63 | Sersi Iēmī idañi issi. | Cersei and Jaime are twins. |
| HighValyrian | 64 | Drīvose aōt bardugon syluti. | Actually we tried to write to you. |
| HighValyrian | 65 | Iōnos sȳz issa. | John is good. |
| HighValyrian | 66 | Mirrori eminna. | I will have whatever. |
| HighValyrian | 67 | Drīvose ropaoty daor. | Actually we are not falling. |
| HighValyrian | 68 | Daor, ñuhon dēmalion issa. | No, it is my throne. |
| HighValyrian | 69 | Bisy zokla issa. | This one is a wolf. |
| HighValyrian | 70 | Riñi valoti annī rhaenis. | The girls find the men’s horses. |
| HighValyrian | 71 | Zȳhor valonqar iā zȳha lēkia iksā? | Are you her younger brother or her older brother? |
| HighValyrian | 72 | Bisi azantyssy issi. | These ones are knights. |
| HighValyrian | 73 | Kastra qintra emi. | We have green turtles. |
| HighValyrian | 74 | Bisy ñuha qȳbranna issa. | This is my cousin. |
| HighValyrian | 75 | Sparos idakēlā? | Whom will you attack? |
| HighValyrian | 76 | Dāria dāri hae mittȳ iotāptetas. | The queen considered the king an idiot. |
| HighValyrian | 77 | Vala sētera izūgas. | The man fears the magic spells. |
| HighValyrian | 78 | Aōhos geltose jomīsā, ñuhus trēsys? | Are you wearing your helmet, my son? |
| HighValyrian | 79 | Konon havon sindīluty daor. | We are not going to buy that bread. |
| HighValyrian | 80 | Skoros aōho mījāeliot liorā? | What are you selling at your pawnshop? |
| HighValyrian | 81 | Varys Mȳrot glaestas. | Varys lived in Myr. |
| HighValyrian | 82 | Ñuhyz dekossa yne ōdris. | My feet are hurting me. |
| HighValyrian | 83 | Vīlībagon gīmīlāt! | You will know how to fight! |
| HighValyrian | 84 | Azantys ñuhe annī urnes. | The knight sees my horse. |
| HighValyrian | 85 | Taoba tolie prūbrī iprattas. | The boy ate another apple. |
| HighValyrian | 86 | Bisy ñuhys raqiros issa. | This one is my friend. |
| HighValyrian | 87 | Ñuhi gelti kostōbi issi. | My helmets are strong. |
| HighValyrian | 88 | Zān taoba olvȳni ipradagon kōttos daor. | Yesterday the boy could not eat much. |
| HighValyrian | 89 | Hontes issa. | It is a bird. |
| HighValyrian | 90 | Riña kese lōtinti ykynas. | The girl is smelling this pie. |
| HighValyrian | 91 | Qaedar lōgor izūgas. | The whale fears the boat. |
| HighValyrian | 92 | Āeksia ñuha drōma sylutesi. | The lords are tasting my eggs. |
| HighValyrian | 93 | Ao ūī dāriot tepā. | You are giving it to the queen. |
| HighValyrian | 94 | Embro sēter ñuho raqiro lōgor jemas. | The ocean’s spell guides my friend’s boat. |
| HighValyrian | 95 | Ñuha kepa jentī rȳbas. | My father hears the leaders. |
| HighValyrian | 96 | Spare azantys ñuhe muñe jorrāelza? | Which knight loves my mother? |
| HighValyrian | 97 | Ābre urnes. | He sees the woman. |
| HighValyrian | 98 | Azantys gevī līritas. | The knight smiled beautifully. |
| HighValyrian | 99 | Zaldrīzes azanto gevivī majaqsa. | The dragon admires the knight’s beauty. |
| HighValyrian | 100 | Kirimvose daor. Toli jorrāelan. | No thank you. I love someone else. |
| Dothraki | 1 | Yer fich anhaán hrakkarés. | You brought me a lion. |
| Dothraki | 2 | Ókki zhílle hrazéf fin állayafa sháfka drogikhoón ánni. | Choose any horse you like from my herd. |
| Dothraki | 3 | Hash me laz ta máe ? | Can she do it? |
| Dothraki | 4 | Atthirár neák khalakkasaán! | Long life to the prince! |
| Dothraki | 5 | Me zíjervo vo rek zhorés! | She showed that heart no mercy! |
| Dothraki | 6 | Hash yer char hakeés? | Did you hear the name? |
| Dothraki | 7 | Me fícha tawakóf máe jinnaán! | He brings his steel here! |
| Dothraki | 8 | Hash sháfka záli memé drívoe? | Do you want him to die? |
| Dothraki | 9 | Réki vácchara máe majín me áchoma. | That will convince him to be respectful. |
| Dothraki | 10 | Ánha zalák áte hazoón! | I want one of those! |
| Dothraki | 11 | Ánha ray vos nakhók! | I’m not finished yet! |
| Dothraki | 12 | Áse máegi ízzi char. | A witch’s words poison the ears. |
| Dothraki | 13 | Vo mawízzi vékho jínne. | There are no rabbits here. |
| Dothraki | 14 | Yer zígeree serj sásha ma láina! | You need a new and stylish vest! |
| Dothraki | 15 | Me véssa atthirár yéri! | It will change your life! |
| Dothraki | 16 | Yazh rhaeshoón ésina! | Spices from different lands! |
| Dothraki | 17 | Ánha ray tih san jáni! | I have seen many dogs! |
| Dothraki | 18 | Ezás loy álegri h’anhaán. | Find some ducks for me. |
| Dothraki | 19 | Jíni dávrae, m’énta fáka. | It’s good for the baby to kick. |
| Dothraki | 20 | Ánha ochomók yeraán kíjinosi. | I will not give you that honor. |
| Dothraki | 21 | Me zígeree mithrát. | He needs to rest. |
| Dothraki | 22 | Kísha ray hezháh chek asshékh. | We’ve ridden far enough today. |
| Dothraki | 23 | Rhéla ánna azzohát máe mra lommayaán. | Help me get him in the tub. |
| Dothraki | 24 | Azhás anhaán haz fes, zhey krísta. | Hand me that carrot, dear. |
| Dothraki | 25 | Jin sérja’thim áffisa. | This vest must be washed. |
| Dothraki | 26 | Yéri vos ánnevo ánna. | You did not leave me. |
| Dothraki | 27 | Yer ayyeyoón lajakoón. | You’ve always been a fighter. |
| Dothraki | 28 | Affín shekh yóla she jímma ma drívoe she títha… | When the sun rises in the west and sets in the east… |
| Dothraki | 29 | Yer ássoo ánna voséchhi! | You do not command me. |
| Dothraki | 30 | Jínne vos gáche vimithrerát. | This is no place to camp. |
| Dothraki | 31 | Ánha vosoón avvós. | I have never been nothing. |
| Dothraki | 32 | Jíni vos jílo. | This is not right. |
| Dothraki | 33 | Vórsa jáda ajjalán! | The fire comes tonight! |
| Dothraki | 34 | Yer ray fich kishaán athohharár! | You’ve brought us destruction! |
| Dothraki | 35 | Yer dévi, zhey krísta, vósma ánha villák. | You’re quick-witted, dear, but I’m wise. |
| Dothraki | 36 | Móri atthasísh oakáh moón. | They killed his soul. |
| Dothraki | 37 | Hash móri vázhi kishaán emralát? | Will they let us in? |
| Dothraki | 38 | Ánha sóqe ákka jin sacchéy éssheyi. | I remove this part of the top. |
| Dothraki | 39 | Yer laz vos véfenari máe vos távi máe vos ívvisi máe. | You can’t pry it or chop it or melt it. |
| Dothraki | 40 | Hash máe vívekhera ma qoyoón ma tolorroón? | Is he made of blood and bone? |
| Dothraki | 41 | Fínne lóshaki? | Where are the guards? |
| Dothraki | 42 | Jíni véna tikh meyér jif ti. | That sounds like something that you would do. |
| Dothraki | 43 | Ma ishísh me átthirarido. | Or maybe it is a dream. |
| Dothraki | 44 | Me’th allayaf mae sekosshi. | He must have been so happy. |
| Dothraki | 45 | Atthirar kishi annevae shorhae. | Our lives have meaning. |
| Dothraki | 46 | Fin yer vijereri? | What do you sell? |
| Dothraki | 47 | Fini thirisir yeri arrek? | How old were you? |
| Dothraki | 48 | Hazi ale khadosoon. | That is more than most have. |
| Dothraki | 49 | Eshna osi vo laini vosso. | The other possibilities are not so pleasant. |
| Dothraki | 50 | Anha zigere yash chosha. | I needed fresh air. |
| Dothraki | 51 | Hash me vallayafa yera tihat mora hezhahhe? | Would you like to see them one day? |
| Dothraki | 52 | Tha yenka onya mazmedha rual fendha yelwa khil. | They shouldn’t have even been allowed to walk our streets. |
| Dothraki | 53 | Nyk skan minty, do jovenne. | I am a soldier, not a politician. |
| Dothraki | 54 | Shkokhé koth pong paza? | How can you trust them? |
| Dothraki | 55 | Verdi ji lysk ilvi qrinuntys zy, do ilvi rageros zy. | We make peace with our enemies, not our friends. |
| Dothraki | 56 | Fin nem olda ki mae? | Who cares about her? |
| Dothraki | 57 | Me azasqana lamekhoon. | She’s paler than milk. |
| Dothraki | 58 | Vos, anha vo zalok vos at rek osoon. | No, I don’t want either of those things. |
| Dothraki | 59 | Anha ray dothra jinne hatif ajjin. | I have been here before. |
| Dothraki | 60 | Ma yer ven toki ven yer shillo mae. | And you were dumb enough to believe him. |
| Dothraki | 61 | Nguwas shonji! | Hold them back! |
| Dothraki | 62 | Awahl utwommani! | Block them off! |
| Dothraki | 63 | Gudniyahl ngugwa yahat todanasa ojigli. | We must be ready before more invaders cross over. |
| Dothraki | 64 | Ngubgukwamanjiyassa, fowanjildiyahl ngugwi. | Either we protect ourselves or we risk losing everything. |
| Dothraki | 65 | Nassa durahahl hafwa. | The threat is too great. |
| Dothraki | 66 | Nyiha guldaniyahl ngugwa nyah. | We must end this now. |
| Dothraki | 67 | Ja já lintot, va jivi kezari. | Or go home, to your families. |
| Dothraki | 68 | Ti morea chek, vosma vitihiri mora. | Treat them well, but keep an eye on them. |
| Dothraki | 69 | Dory umbas. | There are none left. |
| Dothraki | 70 | Inkas hónesko sidri hin bezi. | There are supposed to be more than this. |
| Dothraki | 71 | Tolvyn syri kessa. | Everything will be fine. |
| Dothraki | 72 | Me okeo anni sekosshi. | He is my friend. |
| Dothraki | 73 | Me ohazha memé vo shilo lajat. | Too bad he didn’t know how to fight. |
| Dothraki | 74 | Jorilagon avy sytilibas. | You should go rest. |
| Dothraki | 75 | Ziry ydraon botas. | Let me speak with him. |
| Dothraki | 76 | Uni vali lis va loghor. | All the men have boarded. |
| Dothraki | 77 | Skorkydoso udlilat? | How will you respond? |
| Dothraki | 78 | Ishish me tih leyes. | Maybe she saw a ghost. |
| Dothraki | 79 | Me avvirsae ilek moroa. | It burns their skin. |
| Dothraki | 80 | Eqorasas anna! | Take your hands off me! |
| Dothraki | 81 | Yeri ray esh osoon. | You have made a mistake. |
| Dothraki | 82 | Kisha shilaki yera. | We know who you are. |
| Dothraki | 83 | Yer shillo memé vassila rhaesheseres ma yeroon qisi. | You thought he would conquer the world with you at his side. |
| Dothraki | 84 | Hash yer ray char astosoris mae? | Have you heard the stories about him? |
| Dothraki | 85 | Vosma ei kisha ray tihosh os fin onqotha enossho. | But we all understand the way things are. |
| Dothraki | 86 | Yer ver yomme rhaesheser. | You went out into the world. |
| Dothraki | 87 | Me fasqoyi avezhvenanaz fin laz zali yer, ajjinoon. | It is the best you can hope for, now. |
| Dothraki | 88 | Ma hash rizh yeri norethqoyik? | What if you have a son with red hair? |
| Dothraki | 89 | Anha vovvethak mae ashefasaan. | I’ll throw him in the river. |
| Dothraki | 90 | Hash shafka laz idrie kisha rekkaan akka? | Could you show us the way back? |
| Dothraki | 91 | Mori vo dirgi meqoy jifim ayyoza. | They don’t think the blood should be diluted. |
| Dothraki | 92 | Kijinosi kisha zin hajaki. | That’s how we stay strong. |
| Dothraki | 93 | Hash me jila, jin sen zhavorsi mra qora? | Is it true you have three dragons |
| Dothraki | 94 | Vos yer nem holos anhoon. | Do not betray me. |
| Dothraki | 95 | Me izvena, jin athkessezar az she vaesof. | It is forbidden to carry weapons in the sacred city |
| Dothraki | 96 | Vos at yeroa venoe idrilat mora vosecchi. | None of you is fit to lead them. |
| Dothraki | 97 | Ayos anna jinne. | Wait for me here. |
| Dothraki | 98 | Hash yeri adothrae hrazef ido yomme Havazzhifi Kazga? | Will you ride the wooden horses across the black salt sea? |
| Dothraki | 99 | Hash yeri vaddrivi dozge anni ma khogaroon shiqethi mori majin vohhari okrenegwin mori? | Will you kill my enemies in their iron suits and tear down their stone houses? |
| Dothraki | 100 | Hash yeri m’anhoon, ma jinne m’ayyeyaan?! | Are you with me, now and always?! |

**Supplementary Table 2: Phrases and sentences used in the critical conlang task and their English translations.** (For Dothraki, some phrases and sentences were later discovered to not be in Dothraki. These are highlighted in pink in the table. These phrases and sentences affected n=5 out of the 16 Sentence blocks. We re-modeled the data for Dothraki removing the affected blocks, which did not change the results.)

| **Conlang** | **Number of words per phrase/sentence** | **Phrase/sentence length** | **Number of phrases/sentences per block** | **Run duration** |
| --- | --- | --- | --- | --- |
| Esperanto | Mean = 5.82  SD = 2.00 | Mean = 2.03s  SD = 0.62 | Mean = 6.25  SD = 0.45  Max = 7  Min = 6 | 6:52 (412s) |
| Klingon | Mean = 3.01  SD = 1.45 | Mean = 2.42s  SD = 1.18 | Mean = 6.25  SD = 0.58  Max = 7  Min = 5 | 6:18 (380s) |
| Na’vi | Mean = 5.53  SD = 1.67 | Mean = 3.06s  SD = 0.80 | Mean = 6.25  SD = 0.45  Max = 7  Min = 6 | 7:24 (444s) |
| High Valyrian | Mean = 3.61  SD = 1.05 | Mean = 2.34s  SD = 0.78 | Mean = 6.25  SD = 0.45  Max = 7  Min = 6 | 6:18 (380s) |
| Dothraki | Mean = 4.49  SD = 2.06 | Mean = 2.64s  SD = 1.12 | Mean = 6.25  SD = 0.68  Max = 8  Min = 5 | 6:18 (380s) |

**Supplementary Table 3: Details of the critical conlang experimental paradigm.**

|  | **A1**  “Breakthrough” | **A2**  “Waystage” | **B1**  “Threshold” | **B2**  “Vantage” | **C1**  “Advanced” | **C2**  “Master” |
| --- | --- | --- | --- | --- | --- | --- |
| **Summary of Level** | Ability to understand and use everyday expressions and very basic phrases. | Ability to understand and use common expressions and communicate simple information. | Ability to understand main points of standard input and briefly give reasons and explanations for opinions and plans. | Ability to understand main ideas of complex and abstract topics and interact with a degree of fluency and spontaneity that makes interaction with native speakers possible without strain. | Ability to understand a wide range of demanding, longer clauses, recognize implicit meaning and produce clear, well-structured text on complex subjects. | Ability to understand with ease virtually everything heard or read and to express ideas spontaneously, fluently and with differentiating shades of meaning in most complex situations. |
| **Listening** | I can understand familiar words  and very basic phrases concerning  myself, my family and immediate  concrete surroundings when people speak slowly and clearly. | I can understand phrases and the  highest frequency vocabulary  related to areas of most  immediate personal relevance  (e.g., very basic personal and  Family information, shopping, local  area, employment). I can catch the main point in short, clear,  simple messages and  announcements. | I can understand the main points  of clear standard speech on  familiar matters regularly  encountered in work, school,  leisure, etc. I can understand the main point of many radio or TV programs on current affairs or  topics of personal or professional  interest when the delivery is  relatively slow and clear. | I can understand extended speech  and lectures and follow even  complex lines of argument  provided the topic is reasonably  familiar. I can understand most TV news and current affairs programs. I can understand  the majority of films in standard  dialect. | I can understand extended speech  even when it is not clearly  structured and when relationships  are only implied and not signaled  explicitly. I can understand  television programmes and films without too much effort. | I have no difficulty in understanding  any kind of spoken language,  whether live or broadcast, even  when delivered at fast native  speed, provided I have some time  to get familiar with the accent. |
| **Reading** | I can understand familiar names,  words and very simple sentences,  for example on notices and  posters or in catalogs. | I can read very short, simple  texts. I can find specific,  predictable information in simple everyday material such as  advertisements, prospectuses,  menus and timetables and I can understand short simple personal letters. | I can understand texts that  consist mainly of high frequency  everyday or job-related language. I can understand the description  of events, feelings and wishes in  personal letters. | I can read articles and reports  concerned with contemporary  problems in which the writers adopt particular attitudes or  viewpoints. I can understand  contemporary literary prose | I can understand long and  complex factual and literary texts, appreciating distinctions of style. I can understand specialized  articles and longer technical  instructions, even when they do not relate to my field. | I can read with ease virtually all  forms of the written language,  including abstract, structurally or  linguistically complex texts such as manuals, specialized articles and  literary works. |
| **Spoken**  **Interaction** | I can interact in a simple way  provided the other person is  prepared to repeat or rephrase things at a slower rate of speech and help me formulate what I’m trying to say. I can ask and  answer simple questions in areas of immediate need or on very  familiar topics. | I can communicate in simple and  routine tasks requiring a simple and direct exchange of information on familiar topics and activities. I can handle very short social exchanges, even though I can’t usually understand enough to keep the conversation going myself. | I can deal with most situations  likely to arise whilst traveling in an area where the language is  spoken. I can enter unprepared  into conversation on topics that are familiar, of personal interest  or pertinent to everyday life (e.g., family, hobbies, work, travel and  current events). | I can interact with a degree of fluency and spontaneity that makes regular interaction with native speakers quite possible. I  can take an active part in  discussion in familiar contexts,  accounting for and sustaining my views. | I can express myself fluently and  spontaneously without much  obvious searching for expressions.  I can use language flexibly and  effectively for social and  professional purposes. I can  formulate ideas and opinions with precision and relate my  contribution skilfully to those of other speakers. | I can take part effortlessly in any conversation or discussion and have a good familiarity with idiomatic  expressions and colloquialisms. I  can express myself fluently and convey finer shades of meaning precisely. If I do have a problem I can backtrack and restructure  around the difficulty so smoothly that other people are hardly aware  of it. |
| **Spoken**  **Production** | I can use simple phrases and  sentences to describe where I live and people I know. | I can use a series of phrases and sentences to describe in simple terms my family and other  people, living conditions, my  educational background and my present or most recent job. | I can connect phrases in a simple  way in order to describe  experiences and events, my  dreams, hopes and ambitions. I  can briefly give reasons and  explanations for opinions and  plans. I can narrate a story or relate the plot of a book or film and describe my reactions. | I can present clear, detailed  descriptions on a wide range of  subjects related to my field of  interest. I can explain a viewpoint  on a topical issue giving the  advantages and disadvantages of  various options. | I can present clear, detailed  descriptions of complex subjects  integrating sub-themes,  developing particular points and rounding off with an appropriate  conclusion. | I can present a clear, smoothly flowing  description or argument in a  style appropriate to the context and with an effective logical structure  which helps the recipient to notice and remember significant points. |
| **Writing** | I can write a short, simple  postcard, for example sending  holiday greetings. I can fill in forms with personal details, for example entering my name, nationality and address on a hotel registration form. | I can write short, simple notes and messages. I can write a very  simple personal letter, for example thanking someone for  something. | I can write simple connected text  on topics which are familiar or of personal interest. I can write personal letters describing  experiences and impressions. | I can write clear, detailed text on  a wide range of subjects related  to my interests. I can write an  essay or report, passing on  information or giving reasons in support of or against a particular  point of view. I can write letters highlighting the personal  significance of events and  experiences. | I can express myself in clear,  well-structured text, expressing  points of view at some length. I  can write about complex subjects in a letter, an essay or a report,  underlining what I consider to be the salient issues. I can select a style appropriate to the reader in  mind. | I can write clear, smoothly-flowing  text in an appropriate style. I can write complex letters, reports or  articles which present a case with an effective logical structure which helps the recipient to notice and remember significant points. I can write summaries and reviews of  professional or literary works. |

**Supplementary Table 4**: **Common European Framework of Reference self-assessment grid (Council of Europe, 2001).** Description of the six common reference levels (A1, A2, B1, B2, C1, C2) in the five different domains (listening, reading, spoken interaction, spoken production and writing) that describe varying proficiency levels. Participants were asked to rate their proficiency for each of the domains.

| **Conlang** | **Paradigm** | **Sentences:**  **Mean response** | **Control:**  **Mean response** | **Beta** | **P-Value** |
| --- | --- | --- | --- | --- | --- |
| Esperanto | Conlang | 2.155 | 0.885 | 1.27 | <0.001 |
| Klingon | Conlang | 1.361 | 0.405 | 0.956 | <0.001 |
| Na’vi | Conlang | 2.674 | 1.068 | 1.606 | <0.001 |
| HighValyrian | Conlang | 1.365 | 0.448 | 0.917 | <0.001 |
| Dothraki | Conlang | 1.859 | 0.734 | 1.125 | <0.001 |
| Esperanto | Native language listening | 2.127 | 0.462 | 1.665 | <0.001 |
| Klingon | Native language listening | 1.667 | 0.684 | 0.984 | <0.001 |
| Na’vi | Native language listening | 2.134 | 0.515 | 1.619 | <0.001 |
| HighValyrian | Native language listening | 2.126 | 1.290 | 0.835 | 0.002 |
| Dothraki | Native language listening | 2.120 | 0.639 | 1.481 | <0.001 |
| Esperanto | English reading | 2.071 | 0.675 | 1.396 | <0.001 |
| Klingon | English reading | 2.001 | 0.857 | 1.144 | <0.001 |
| Na’vi | English reading | 2.675 | 0.929 | 1.746 | <0.001 |
| HighValyrian | English reading | 1.731 | 0.604 | 1.126 | <0.001 |
| Dothraki | English reading | 2.373 | 0.788 | 1.586 | <0.001 |

**Supplementary Table 5: Results of the linear models fit separately for each of the conlang groups (i.e., Esperanto, Klingon, Na’vi, High Valyrian, Dothraki) examining the responses of the language areas to the Sentences and Control conditions of the language localizers and the critical conlang task.** The Paradigm, Condition Response column refers to the average BOLD response across the language areas of the condition listed in the Paradigm, Condition column. The Beta and p-value columns correspond to values resulting from fitting the following linear model for each conlang group (with the sentence condition being the reference): *EffectSize ~ Condition + (1|Participant) + (1|ROI)***.**

| **Conlang** | **Conlang Sentence Response** | **Paradigm, Condition** | **Paradigm, Condition Response** | **Beta** | **P-value** |
| --- | --- | --- | --- | --- | --- |
| Esperanto | 2.155 | Spatial Working Memory – Hard | -0.424 | -2.62 | <0.001 |
| Esperanto | 2.155 | Spatial Working Memory – Easy | -0.228 | -2.437 | <0.001 |
| Esperanto | 2.155 | Theory of Mind – Mental | -0.038 | -2.249 | <0.001 |
| Esperanto | 2.155 | Theory of Mind – Physical | 0.323 | -1.872 | <0.001 |
| Klingon | 1.361 | Spatial Working Memory – Hard | -0.443 | -1.761 | <0.001 |
| Klingon | 1.361 | Spatial Working Memory – Easy | -0.364 | -1.69 | <0.001 |
| Klingon | 1.361 | Theory of Mind – Mental | 0.01 | -1.329 | <0.001 |
| Klingon | 1.361 | Theory of Mind – Physical | 0.019 | -1.264 | <0.001 |
| Na’vi | 2.674 | Spatial Working Memory – Hard | -0.403 | -3.095 | <0.001 |
| Na’vi | 2.674 | Spatial Working Memory – Easy | -0.196 | -2.887 | <0.001 |
| Na’vi | 2.674 | Theory of Mind – Mental | 0.083 | -2.623 | <0.001 |
| Na’vi | 2.674 | Theory of Mind – Physical | -0.345 | -3.02 | <0.001 |
| HighValyrian | 1.365 | Spatial Working Memory – Hard | -0.295 | -1.682 | <0.001 |
| HighValyrian | 1.365 | Spatial Working Memory – Easy | -0.297 | -1.7 | <0.001 |
| HighValyrian | 1.365 | Theory of Mind – Mental | -0.221 | 1.652 | <0.001 |
| HighValyrian | 1.365 | Theory of Mind – Physical | -0.088 | -1.544 | <0.001 |
| Dothraki | 1.859 | Spatial Working Memory – Hard | -0.017 | -1.976 | <0.001 |
| Dothraki | 1.859 | Spatial Working Memory – Easy | -0.046 | -2.017 | <0.001 |
| Dothraki | 1.859 | Theory of Mind – Mental | -0.711 | -2.645 | <0.001 |
| Dothraki | 1.859 | Theory of Mind – Physical | -0.632 | -2.557 | <0.001 |

**Supplementary Table 6**: **Results of the linear models fit separately for each of the conlang groups (i.e., Esperanto, Klingon, Na’vi, High Valyrian, Dothraki) examining the responses of the language areas to the Sentences condition of the conlang task and each of the four non-linguistic conditions.** The Paradigm, Condition Response column refers to the average BOLD response across the language areas of the condition listed in the Paradigm, Condition column. The Beta and p-value columns correspond to values resulting from fitting the following linear model for each conlang group (with the sentence condition being the reference): *EffectSize ~ Condition + (1|Participant) + (1|ROI)***.**

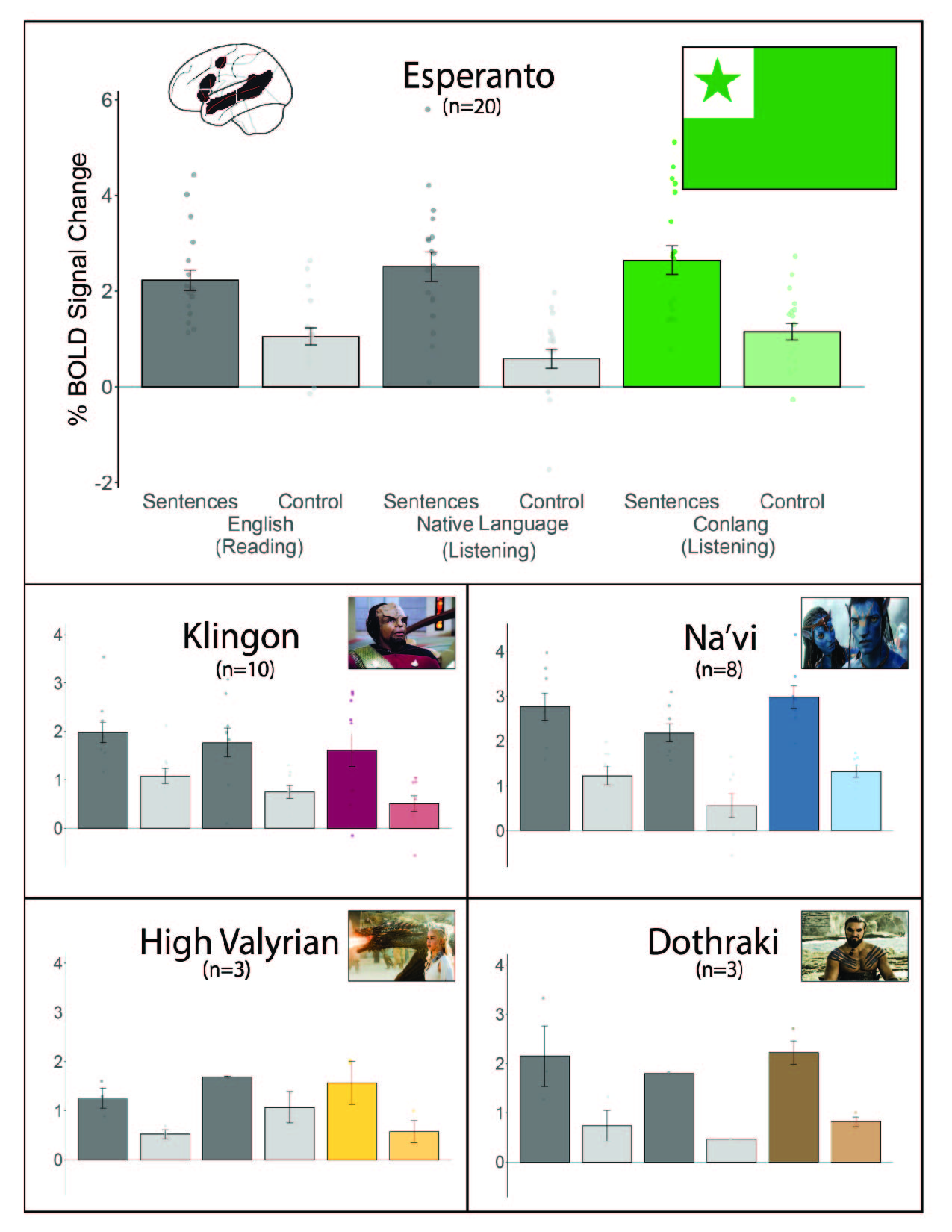

**Supplementary Figure 1: The language areas (defined by the Sentence>Control contrast in the native language listening task; cf. English) show a strong response during the processing of both natural and constructed languages.** Percent BOLD signal change across the five language fROIs for the reading-based and the listening-based versions of the language localizer (dark grey bars = sentences, light grey bars = control condition) and the critical conlang comprehension task, where participants listened to sentences in the constructed language (colored bars) and to control, acoustically degraded versions of those sentences (lighter versions of the colored bars). The language network is defined by the Sentences>Control condition of the listening language localizer in the language-dominant hemisphere (left hemisphere for 36 of the 38 participants). To quantify the similarities between the language network defined by the different tasks (i.e., as defined by the Sentences>Control contrast in the native language listening task and by the Sentences>Control contrast in the reading task), a voxel-wise correlation within the language areas in the 36 native English speaking participants was computed; the two were correlated (Fisher-transformed) at r=0.750 (SD=0.357). Dots represent individual participants; error bars represent standard errors of the mean by participant.

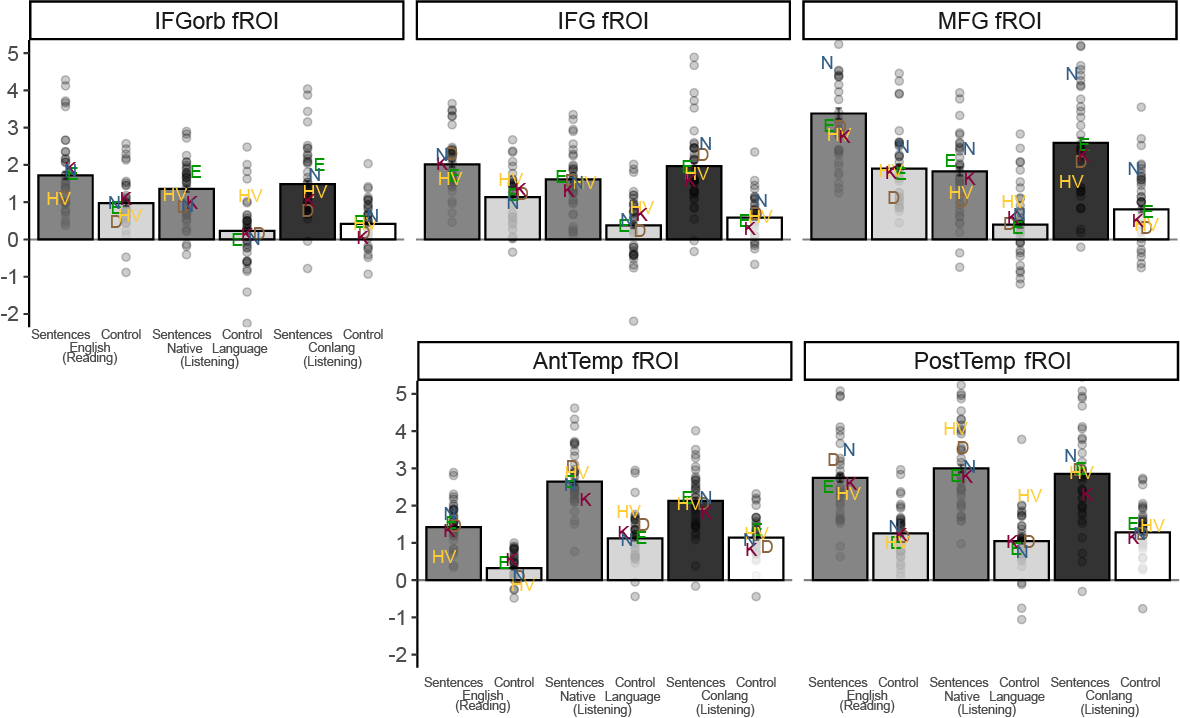

**Supplementary Figure 2: Each of the fROIs in the language network shows a strong response during the processing of both natural and constructed languages.** Percent BOLD signal change in the five language fROIs for the reading-based and the listening-based versions of the language localizer (dark grey bars = sentences, light grey bars = control condition) and the critical conlang comprehension task, where participants listened to sentences in the constructed language (black bars) and to control, acoustically degraded versions of those sentences (white bars). The language network is defined by the Sentences>Control condition of the English reading localizer in the language-dominant hemisphere (left hemisphere for 36 of the 38 participants). Dots represent individual participants; letters show the average response for each conlang group (E: Esperanto; K: Klingon; N: Na’vi; HV: High Valyrian; D: Dothraki); error bars represent standard errors of the mean by participant.

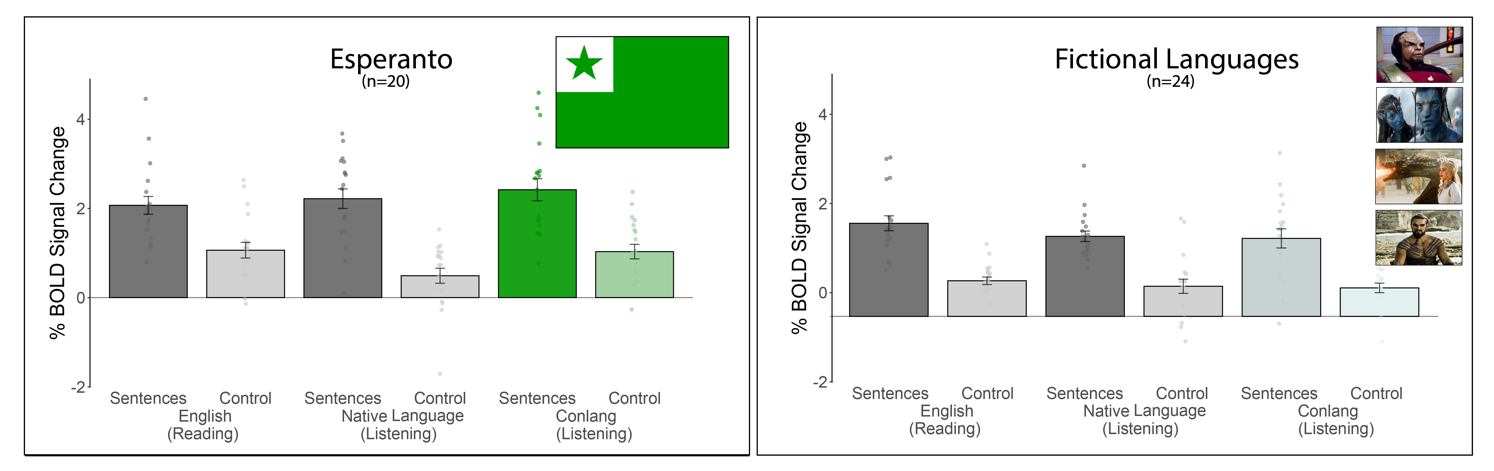

**Supplementary Figure 3: The responses to Esperanto and four fictional conlangs in language regions are comparable.** Percent BOLD signal change across the five language fROIs for the reading-based and the listening-based versions of the language localizer (dark grey bars = sentences, light grey bars = control condition) and the critical conlang comprehension task, where participants listened to sentences in the constructed language (colored bars) and to control, acoustically degraded versions of those sentences (lighter versions of the colored bars). Dots represent individual participants; error bars represent standard errors of the mean by participant. Results from a linear mixed effects model where Conlang was entered as a fixed effect, and participant and ROI were entered as random effects, did not yield any statistical significance between Esperanto and Klingon (p=0.48), Na’vi (p=0.25), High Valyrian (p=0.35) and Dothraki (p=0.21).
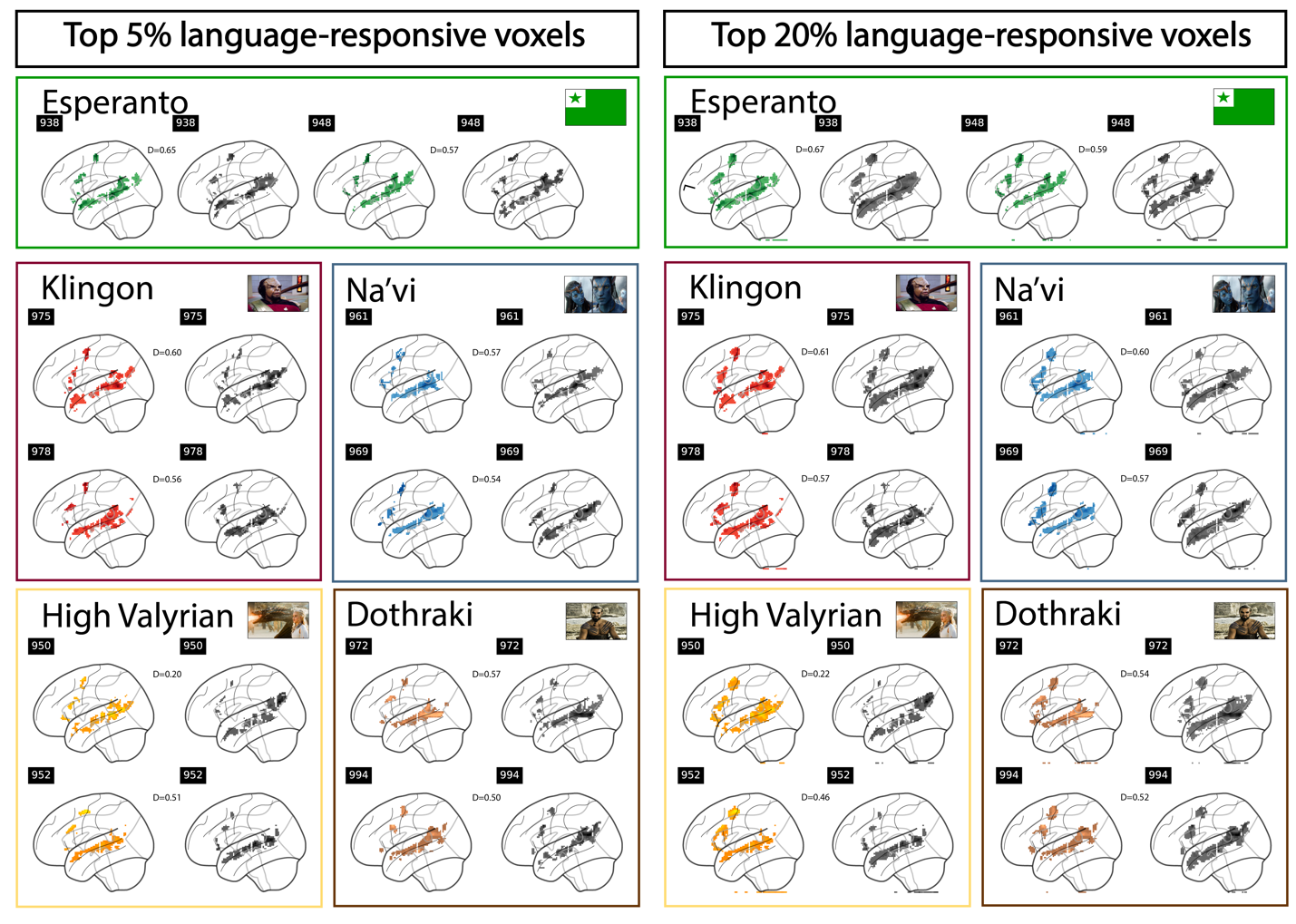

**Supplementary Figure 4: The amount of overlap (Dice coefficient) is robust to the percentage of voxels taken to define language-responsive voxels.** Binarized versions of the activation maps for the Sentences>Control contrast for the participant’s native language and the relevant conlang where—restricting the analysis to the LH language parcels—voxels that belonged to the set of 5% (left) and 20% (right) of most responsive voxels were assigned a value of 1 and voxels that did not were assigned a value of 0. We then computed the Dice coefficient (Rombouts et al., 1997), which ranges from 0 (no overlapping voxels) to 1 (all voxels overlap) between a) the native language map and the conlang map within each participant (n=38 pairs), and b) the native language maps between each pair of participants (n=703 pairs). We found that the sets of most responsive voxels are more similar (show greater overlap) within an individual between a natural and a constructed language than between individuals for a natural language both when taking the top 5% of voxels (0.50 vs. 0.21; β=0.28, p<0.001) and when taking the top 20% of voxels. voxels (0.53 vs. 0.35; β=0.18, p<0.001). Moreover—across participants—the degree of proficiency in the conlang was associated with the degree of similarity between the conlang activations and native language activations (r=0.395, p<0.01).

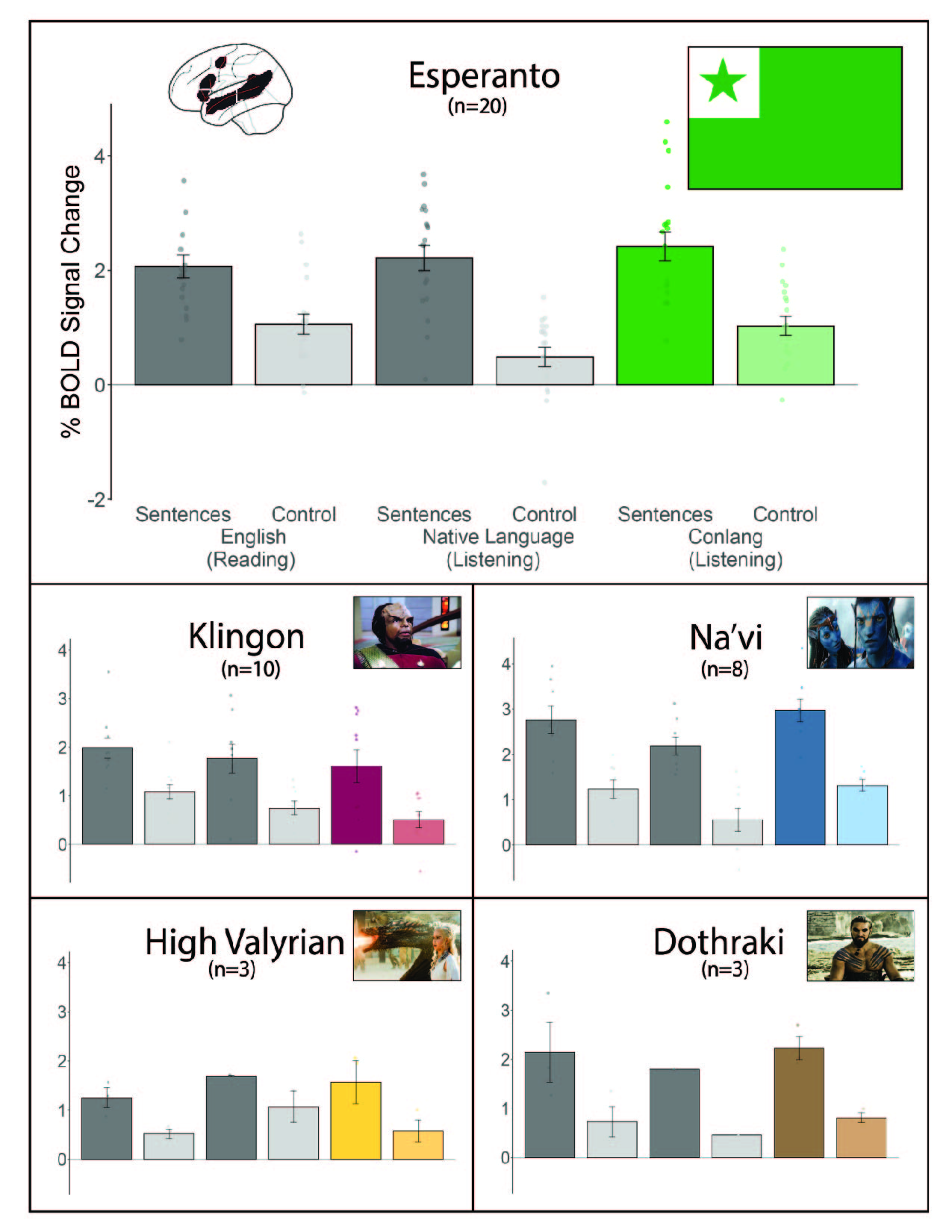

**Supplementary Figure 5: The language areas (in the left hemisphere; cf. the language-dominant hemisphere) show a strong response during the processing of both natural and constructed languages.** Percent BOLD signal change across the five language fROIs for the reading-based and the listening-based versions of the language localizer (dark grey bars = sentences, light grey bars = control condition) and the critical conlang comprehension task, where participants listened to sentences in the constructed language (colored bars) and to control, acoustically degraded versions of those sentences (lighter versions of the colored bars). The language network is defined by the Sentences>Control condition of the English reading localizer in the left hemisphere (cf. Figure 1 for responses in the dominant hemisphere, as 2 of the 38 participants had right-lateralized language networks). Dots represent individual participants; error bars represent standard errors of the mean by participant.

| **A. Whole-brain maps for the random-effects group analysis** | |
| --- | --- |
| **Multiple Demand Network**  (Hard > Easy Contrast) | **Theory of Mind Network**  (Mental > Physical Contrast) |
| 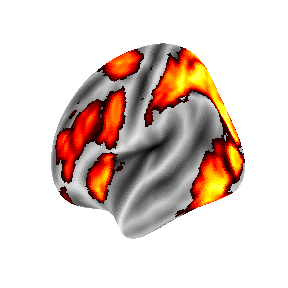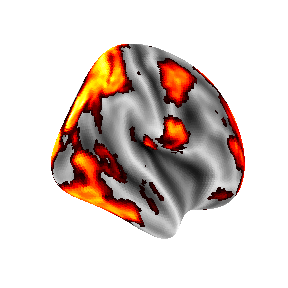  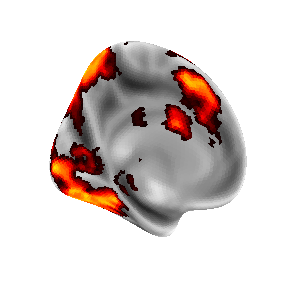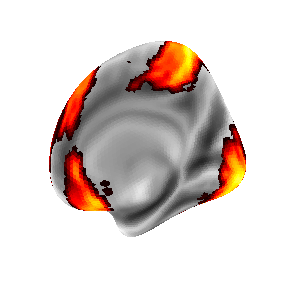 | 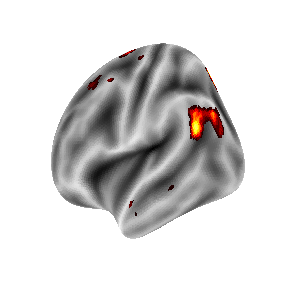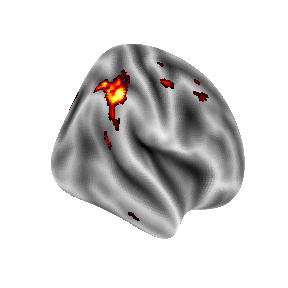  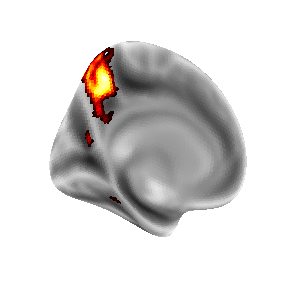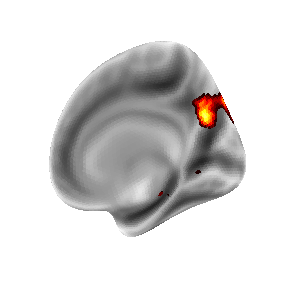 |
| **B. Percent BOLD signal change across the Multiple Demand and Theory of Mind fROIs**  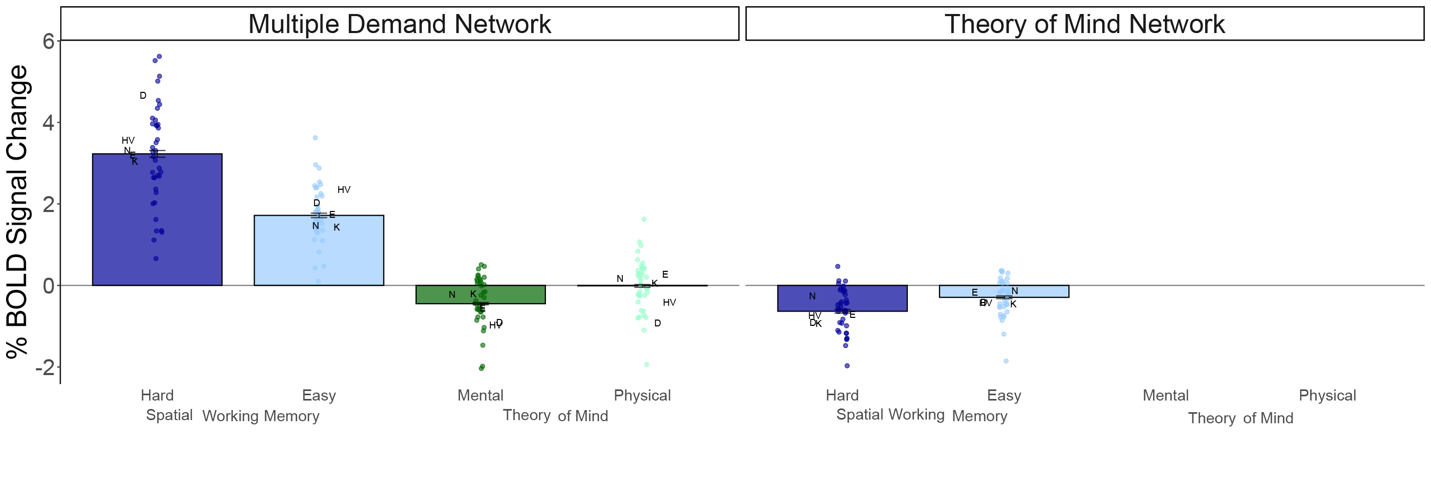 | |

**Supplementary Figure 6:** **Whole-brain group-level activation maps for non-linguistic tasks and** **responses to non-linguistic tasks in the Multiple Demand and the Theory of Mind fROIs. A.** Whole-brain activation maps for the random-effects group analyses for the *Hard>*Easy contrast of the spatial working memory task, and the *Mental>Physical* contrast of the Theory of Mind task. The maps are shown thresholded at p<0.001 uncorrected level. For each task, the top two images correspond to the LH and RH lateral views, and the bottom two images—to the LH and RH medial views. The maps for both tasks show the expected topography (e.g., Duncan, 2013; Jacoby et al., 2016). **B.** Percent BOLD signal change across the Multiple Demand (MD) and Theory of Mind (ToM) fROIs for the conditions of the spatial working memory task and the naturalistic ToM task. The MD network is defined by the *Hard>Easy* contrast of the spatial working memory task (Fedorenko et al., 2013); the ToM network is defined by the *mental>physical* contrast for the naturalistic ToM task (Jacoby et al., 2016). (Given that a) an across-runs cross-validation procedure is used to ensure independence between the data used to define the fROIs vs. to examine the response magnitudes, and b) the naturalistic ToM task consists of a single run, responses to the ToM task conditions cannot be estimated in the ToM network.) Dots represent individual participants; letters show the average response for each conlang group (E: Esperanto; K: Klingon; N: Na’vi; HV: High Valyrian; D: Dothraki); error bars represent standard errors of the mean by participant.

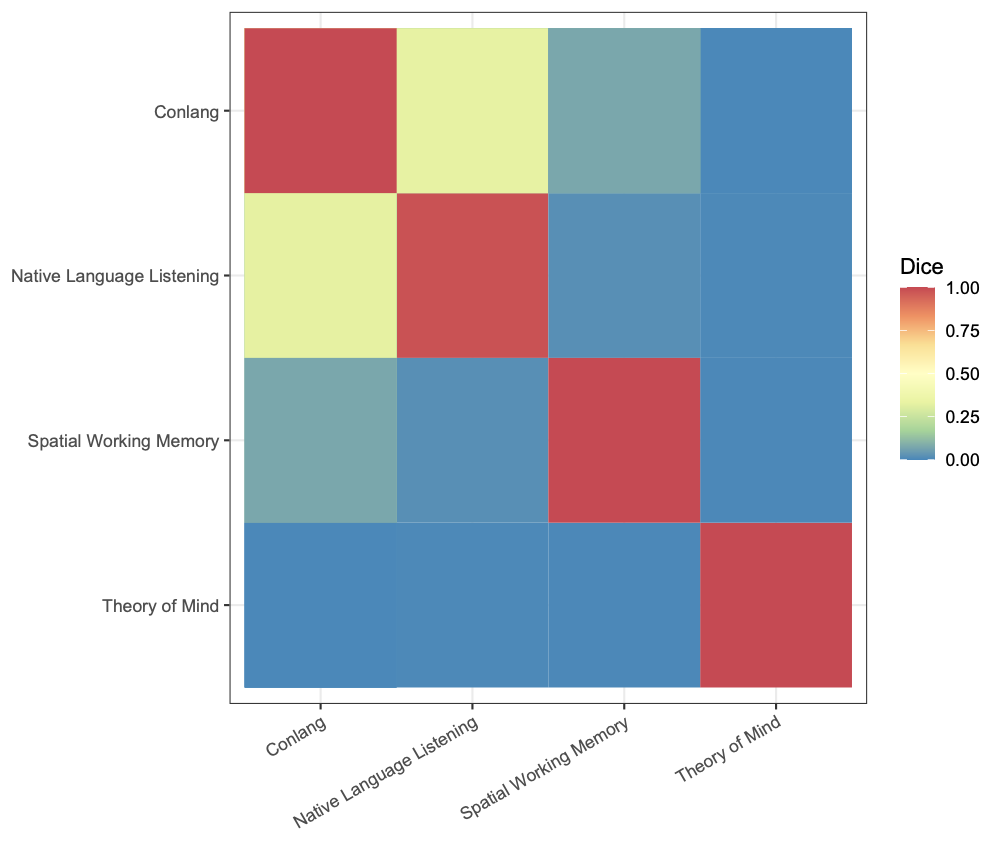

**Supplementary Figure 7: Whole-brain individual-level activation overlap between conlang responses and responses to the language localizer vs. non-linguistic tasks.** The Dice coefficient (Rombouts et al., 1997) was calculated between individual-level activation maps for four contrasts (the *Sentences>Control* contrast for the conlang task and the native language listening task, the *Hard>Easy* contrast for the spatial working memory task, and the *mental>physical* contrast for the Theory of Mind (ToM) task). For each individual and for each contrast, the whole-brain maps were thresholded at p<0.001 uncorrected level and binarized such that significant voxels were assigned a value of 1 and non-significant voxels—a value of 0. Then, in each individual participant for each pair of contrasts, we calculated the number of overlapping voxels, multiplied this value by 2, and then divided it by the total number of significant voxels across the two images (i.e., Dice Similarity Coefficient(A,B) = 2(A∩B)/(A+B)). The resulting values range from 0 (no overlapping voxels) to 1 (all voxels overlap). This procedure resulted in a total of 128 comparisons (given that 5 of the 38 participants spoke two conlangs, and one participant did not complete the ToM task, there were a total of 43 comparisons for the conlang-native language listening and conlang-spatial working memory and 42 comparisons for the conlang-ToM). These comparisons were then averaged across participants for each contrast pair. Activations for the conlang task showed significantly greater overlap with the activations for the native language listening task than with either the spatial working memory task (0.27 vs. 0.06, β = 0.20, p<0.001) or the ToM task (0.27 vs 0.001; β = 0.27, p<0.001).
